# TAUS: Target-Age Unified Survival. Survival analysis without assuming proportional hazards or parameterising the survival function

**DOI:** 10.64898/2026.04.15.718114

**Authors:** Iván Casas Gomez-Uribarri, Simon A. Babayan, Fredros Okumu, Francesco Baldini, Mauro Pazmiño Betancourth

**Affiliations:** School of Biodiversity, One Health & Veterinary Medicine, University of Glasgow, Glasgow, UK; Ifakara Health Institute, Ifakara, Morogoro region, Tanzania

**Keywords:** Survival analysis, Cox Proportional Hazards, Parametric survival models, R package, TAUS, Kaplan-Meier, Conditional probability, Demography

## Abstract

1. Standard survival analysis methods often rely on the assumption of proportional hazards (PH) or parameterisations of the survival function that might not be appropriate for wild populations.
2. To enable survival analysis without these modelling constraints, we developed an approach that combines the Kaplan-Meier estimator with conditional probability theory to compute age-specific probabilities of survival up to some target age of choice *τ*. Marginalising this probability over the age distribution of the population yields *O*_*τ*_, the probability that a randomly sampled individual of unknown age will outlive the target age *τ*. Notably, the value for *τ* is set by the analyst for each group independently, which allows accounting for differences in pace of life across populations.
3. We tested its application using a simulation study and two real-world datasets, and compared its performance against that of Cox PH and parametric survival models. The PH assumption was violated in the three examples, rendering the Cox PH models inappropriate. Parametric models offered a better alternative, but the best parametric fit missed at least some key survival patterns in all examples. The TAUS model provided a valid description of survival patterns in all cases. Its richer output also allowed finer analysis of survival differences between populations.
4. The `TAUS` model is also available as an R package (https://github.com/casasgomezuribarri/TAUS). This new approach to survival analysis without PH or parametric assumptions allows the comparison of survival probabilities across populations with different age structures and rates of pace of life. This makes it suitable for a wide range of ecological applications, including in population viability analysis, epidemiology, or life-history theory

## INTRODUCTION

Survival analysis is the statistical study of time-to-event data, where the outcome variable is the time between two observable events. In practice, the first event is usually the start of the study or process, and the outcome is the time until the event of interest occurs. It is the convention to refer to this event of interest as *death*, although its nature is not restricted to biological death. The terminology is coherent with this convention: *birth* refers to the start of the process under study, *age* is the time since birth, and *lifespan* is the typical age at death. This semantic convention will be applied throughout this manuscript for simplicity, but it should be noted that survival analysis is not restricted to the analysis of biological survival.

Understanding how survival patterns change across populations, time, and environmental conditions is central to many questions in ecology and evolutionary biology, with direct applications in conservation, epidemiology, or life-history theory [Nur et al., 2004, Scherm and Ojiambo, 2007, Selvin, 2010, Heppell, 1998]. However, the most common methods for survival analysis—Cox proportional hazards (PH) and parametric survival models—are often inadequate for use in ecological contexts.

Cox PH models investigate the effect of one or more explanatory variables on survival by estimating the ratio of the hazards of different groups, where hazard is defined as the age-specific immediate risk of death, given survival to that age [Cox, 1972]. A central assumption of the model is that this ratio is constant over time, but this is often violated in practice [Kuitunen et al., 2021, Dunkler et al., 2018, Kleinbaum and Klein, 2012]. Methods to relax this limitation include stratifying the data by the variable that violates the assumption [Kleinbaum and Klein, 2012], partitioning the timeline into smaller intervals for which the hazards ratio is assumed to be constant [Kuitunen et al., 2021], or applying weights to different timepoints to estimate the *average* hazard ratio [Dunkler et al., 2018]. Despite these extensions, Cox PH models remain limited to relative descriptions of survivorship —providing no absolute estimates of individual hazards— and inherently unable to describe how covariate effects change over time. Indeed, Cox himself noted the difficulty of giving a clear physical or biological interpretation of the model’s output, and later expressed a preference for parametric approaches when analysing censored survival data [Cox, 1997, Cox and Reid, 1994].

Parametric survival models, the second most common approach to survival analysis, assume that survival times follow some known probability distribution, and then estimate its parameters as a function of the explanatory variables [Crowther and Lambert, 2014]. Unlike the Cox PH model, they can produce absolute survival estimates and detect effects that are time-dependent [Crowther and Lambert, 2013]. However, they rely on the good fit of the chosen distribution for the baseline hazard function, and consequently inherit its structural constraints. These include restrictions on modality, hazard shape, parameterisation, and on distributional properties such as symmetry or tail behaviour. It has been shown that, in this context, model misspecification can cause bias of clinically-relevant magnitude in medical applications of survival analysis [Bellera et al., 2010, Nardi and Schemper, 2003, Gasparini et al., 2018]. Parametric survival models are also limited in their ability to compare survival dynamics across populations that follow different distributions, even when their independent fits are adequate.

Here, we present a novel approach to survival analysis that does not parameterise the hazard function or assume proportional hazards. Our model produces absolute survival estimates, which come in the form of the predicted probability of outliving some age *τ*. More specifically, it quantifies the probability that a *randomly sampled individual from a population of known (or estimated) age distribution, but whose current age is unknown* will live to be at least age *τ*. This metric is particularly useful in ecology, or more broadly in contexts where:

- Estimating the age structure of a population may be more feasible and cost-effective than age-grading every individual, which is often the case when working with wild populations [Müller et al., 2004, Johnson et al., 2020]. For example: *Given that the age of each fish is unknown but the age distribution in each loch can be estimated, what is the difference between loch A and loch B in the probability that a randomly sampled fish will survive beyond age τ?*
- The research question concerns survival beyond a target age that may differ between populations, as in population viability analysis [Eberhardt, 2002], epidemiology [Ohm et al., 2018], or when age is quantified via biophysical measures rather than time [Ludwig and Smoke, 1980, Jackson et al., 2003]. For example: *In which of two allopatric populations is a randomly sampled individual more likely to survive beyond sexual maturity, considering that sexual maturity is reached at a different age in each population?*
- The research question is conditional on the event occurring within (or outside) a certain age window. For example, since menopause before age 45 has been associated with all-cause mortality [Muka et al., 2016], one might ask: *What is the probability that a female from population A experiences menopause before age 45, compared with a female from population B?*.

We use a simulation study and two real-world datasets to illustrate how to implement this model and how to interpret its output. We also compare its statistical robustness and ecological value against those of traditional approaches to survival analysis (i.e. Cox PH and parametric survival models).

### Survival analysis: an overview

Survival analysis is relevant to many fields, including biology, medicine, engineering, and others. In practice, survival analysis involves the study of the probability of death, and of how this probability changes over time or as a function of some explanatory variables [Emmert-Streib and Dehmer, 2019]. This is usually modelled as follows:

Let *T* be a nonnegative random variable denoting time until death. Its density function *f* (*t*) is defined by:

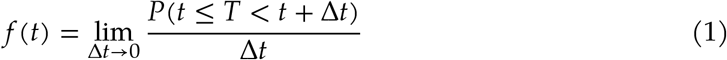

The cumulative distribution function *F*(*t*) shows the probability that death happens before or exactly at time *t*:

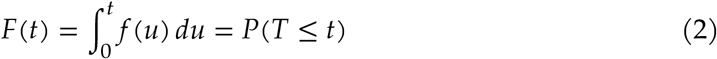

From here, one can derive the survival function *S*(*t*) which depicts the probability that death will happen after time *t* (i.e. probability of survival until *t*):

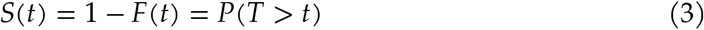

Lastly, we can derive the hazard function *h*(*t*), which illustrates the probability of immediate death at time *t* conditional on survival up to that time:

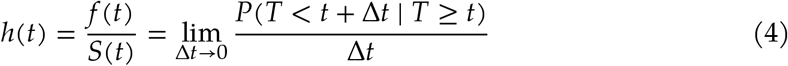

In practice, the true survival function of a population is unknown and must be estimated from sample data, which are often censored. For censored individuals, only partial information is available: their age at death is unknown, but it is known that death did not occur within a specific time interval. Statistical methods allow estimation of a population’s survival function from such censored data. The Kaplan-Meier (KM) estimator provides a nonparametric estimate of *S*(*t*) as a step function, properly accounting for censored observations. However, KM is limited to estimating survival curves within individual populations, and cannot be used to evaluate the effects of covariates on survival. For this purpose, the most common tools are the Cox and parametric approaches described above. Here, we present an alternative.

## METHODS

This section outlines the theoretical framework of the model. First, a formula is derived for the conditional survival function, which describes the age-specific probability of outliving some age *τ*. This function is then marginalised over the age distribution of the population to yield the probability that a randomly sampled individual of unknown age will outlive age *τ* (*O*_*τ*_). This is followed by a comment on common challenges when estimating the age distribution from data. We then propose a statistical framework for the comparison of two values of *O*_*τ*_. Finally, we enumerate the assumptions made by this model, and summarise its inputs and outputs.

At the end of this section, we describe the (simulated and real-world) datasets used to demonstrate the model’s application in the Results section (Section 4).

### Conditional survival

We derive a formula for the probability that an individual of age *t* will outlive age *τ*. Let *T* denote the time at death (a nonnegative random variable). Applying Bayes’ theorem, we have:

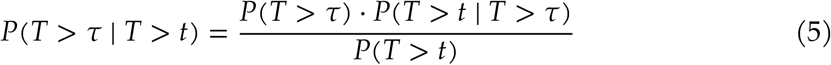

Now, applying Equation 3, we arrive at the following:

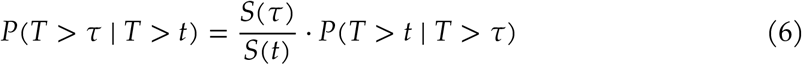

where *S*(*t*) and *S*(*τ*) can be estimated from data using a standard KM model. We can structure the calculation of *P*(*T* > *t* ∣ *T* > *τ*) in two logical cases:

### The case when *t* ≤ *τ*

In this case, the individual’s current age *t* is less than the age of interest *τ*. If an individual has outlived *τ*, it is certain that it has outlived *t*. This leads to the following conditional survival formula, substituting *P*(*T* > *t* ∣ *T* > *τ*) = 1 into Equation 6:

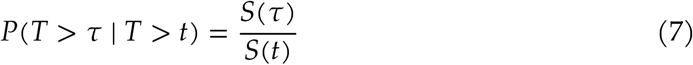

### The case when *t* ≥ *τ*

Following the same logic, when *t* ≥ *τ, P*(*T* > *τ* ∣ *T* > *t*) = 1 (i.e. it *has happened already*).

Therefore, the probability of survival to age *τ* conditional on survival to age *t* can be formulated as follows:

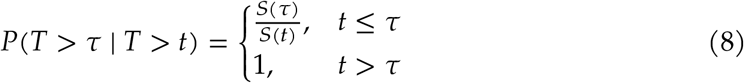

It should be noted that the conditional survival space is defined for all values of *t* and *τ* that are smaller than the largest age recorded in the study.

The conditional survival function *P*(*T* > *τ* ∣ *T* > *t*) (Equation 8) describes the probability that an individual of age *t* will outlive age *τ*. This contrasts with the traditional survival function *S*(*t*), which represents at-birth survival probabilities to age *t*. Indeed, if we set *t* = 0, we get *P*(*T* > *τ* ∣ *T* > 0) = *P*(*T* > *τ*) = *S*(*τ*) (see Equation 3).

### Probability of outliving age *τ* (*O*_*τ*_)

To compare conditional survival across groups, variation must be considered along two dimensions: the age of the individual (*t*), and the age beyond which survival is of interest (*τ*). This makes the comparison of conditional survival more complex than the comparison of other traditional survival functions, which vary only along one dimension.

In order to simplify this comparison for practical applications, we propose that a choice of (one or a few values for) *τ* be made by the analyst. We also propose that the conditional survival *P*(*T* > *τ* ∣ *T* > *t*) be collapsed over the *t* dimension by considering the distribution of *t* (i.e. the age structure of the population). Hence, *O*_*τ*_ can be defined as the probability that a randomly sampled individual of unknown age will outlive age *τ*. By the law of total probability, this is the expectation of *P*(*T* > *τ* ∣ *T* > *t*) over the age distribution: *O*_*τ*_ = 𝔼_A(t)_[*P*(*T* > *τ* ∣ *T* > *t*)]. Treating time discretely, this metric can be calculated as follows:

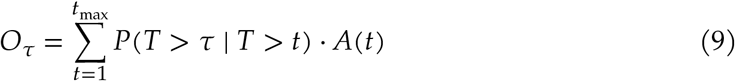

where A(*t*) is the proportion of individuals in a population with current age *t*, also known as the age distribution of the population. Figure 1 illustrates how this metric is calculated for the arbitrary case of *τ* = 15: *O*_*τ*_ equals the area under the red curve, which is the *t*-wise product of *P*(*T* > *τ* ∣ *T* > *t*) (black line) and A(*t*) (blue bars). Section 3.3 discusses different approaches to estimating A(*t*), and Section 3.5 illustrates how uncertainty is propagated to provide a confidence interval around *O*_*τ*_.

**Fig. 1:**
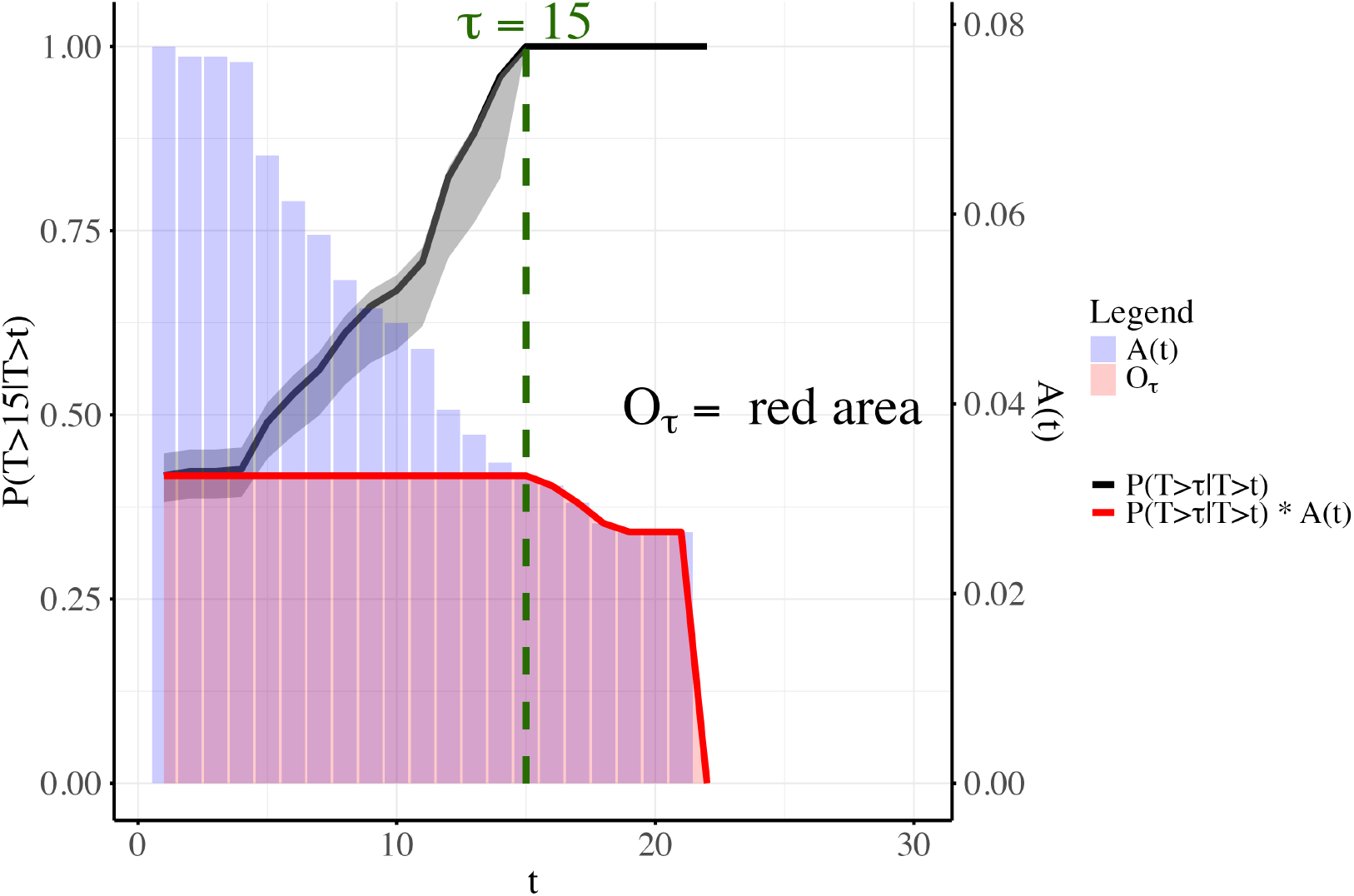
Estimation of *O*_*τ*_ for the arbitrary case of *τ* = 15 (green dashed line). It shows the conditional survival *P*(*T* > *τ* ∣ *T* > *t*) (black line, 95% CI in grey), the age distribution *A*(*t*) (blue histogram), the t-wise evaluation of *P*(*T* > *τ* ∣ *T* > *t*) · *A*(*t*) (red line), and the value of 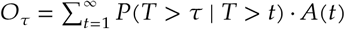 (red area; in practice the sum is over *t* = 1, …, *t*_max_).

Computing *O*_*τ*_ requires a choice of *τ* to be made (in Figure 1, the arbitrary choice was *τ* = 15). Alternatively, *O*_*τ*_ can be plotted as a function of *τ*, producing a curve that we have called *O*(*τ*). This curve—as opposed to the survival function *S*(*t*), which illustrates the probability *at birth* of surviving beyond age *t*—illustrates the probability *at the age of sampling* of surviving beyond age *τ*, where the age at sampling is estimated as an A(*t*)-weighted average of all sampleable ages.

### Estimating A(*t*)

In order to simulate the process of randomly sampling an individual from a population, an estimation of its age distribution A(*t*) is required. One way of achieving this is from survival and fertixlity data. If the population is stable (i.e. constant age-specific birth and death rates, and no migration), A(*t*) can be shown to converge to a stable equilibrium [Lotka, 1937]. This time-invariant A(*t*) can be calculated by constructing the Leslie matrix (which summarises age-specific fertility and mortality) and projecting it forward in time [Caswell, 2001], or by calculating the eigenvector of its dominant eigenvalue [Howard, 2026]. A special case of stable population is a stationary one (i.e. no growth and no migration), which has A(*t*) equal to its normalised survival function *S*(*t*) [Inaba, 2017].

Wild populations are rarely stable, however, due to density-dependent and stochastic environmental processes. These can be modelled by specifying Leslie matrices that are themselves functions of time or of population size [Caswell, 2001]. The age distribution of a non-stable population is not guaranteed to converge to a stable equilibrium: convergence to periodic or chaotic attractors is also possible [Caswell, 2001, Chu, 2015].

Since the objective of estimating A(*t*) is to simulate the process of randomly sampling an individual of age *t*, an alternative is to simulate this process directly from empirical data. If the ages of a random sample of individuals from the population can be determined, then A(*t*) can be approximated as the normalised distribution of the age frequencies of this sample. Additionally, methods have also been proposed to estimate A(*t*) from capture-recapture data of wild individuals when the age at first capture is unknown [Müller et al., 2004, Müller et al., 2007].

### Statistical analysis

To compare two values of *O*_*τ*_, they can be modelled as Beta-distributed random variables, and the distance between the distributions can be calculated using the Kolmogorov-Smirnov (KS) test. Then, a p-value can be computed through bootstrapping to assess the statistical significance of the KS distance between the two distributions. This section proposes a statistical framework for the comparison of two values of *O*_*τ*_.

The Beta distribution is a continuous probability distribution defined in the interval [0, 1], and it is commonly used to model probabilities. It has two shape parameters, *α* and *β*. The mean and variance of a Beta-distributed random variable *X* ~ Beta(*α, β*) are given by:

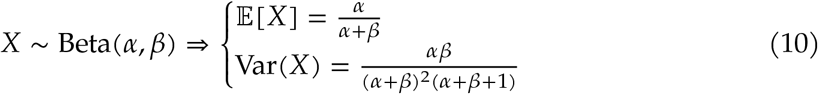

Following this definition, *O*_*τ*_ can be modelled by setting 𝔼(*X*) ≈ *O*_*τ*_ and Var(*X*) ≈ (*U*− *L*)^2^, where *U* and *L* are the upper and lower limits of the 95% CI for *O*_*τ*_, respectively (see Section 3.5 for the calculation of these). Writing *μ* = 𝔼[*X*] and *σ*^2^ = Var(*X*), Equation 10 gives *μ* = *α*/(*α* + *β*) and *σ*^2^ = *αβ*/((*α* + *β*)^2^(*α* + *β* + 1)). Solving for *α* and *β* in terms of *μ* and *σ*^2^ yields 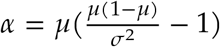 and 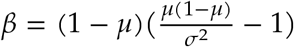. Substituting *μ* = *O*_τ_ and *σ*^2^ = (*U* − *L*)^2^ gives:

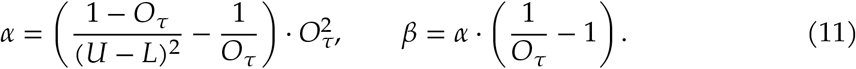

Once the Beta distributions are defined, the KS distance between them is the maximum vertical difference between their two cumulative functions:

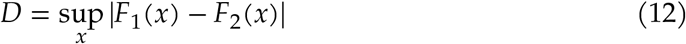

where *F*_1_(*x*) and *F*_2_(*x*) are the cumulative distribution functions of the two Beta distributions being compared, as evaluated at *x*.

In order to test the statistical significance of this distance *D*, we first fit a null Beta distribution to both values of *O*_*τ*_ under comparison. We estimate the null distribution by averaging the *α* and *β* parameters of the two independently-fitted Beta distributions:

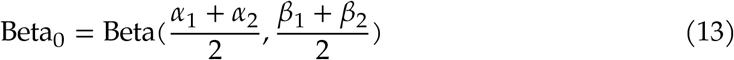

where *α*_1_, *β*_1_, *α*_2_ and *β*_2_ are the parameters of the fitted Beta distributions of the two *O*_*τ*_ values under comparison. We then draw pairs of samples from Beta_0_ and calculate their KS distance using the R function `stats::ks.test`. We repeat this process many times and store the results (i.e. bootstrap). A p-value is then computed as the proportion of these KS distances that are larger than the numerically computed *D* (Equation 12).

This number is the probability of obtaining a KS distance *D* or larger under the null hypothesis that the two *O*_*τ*_ values come from the same Beta distribution (i.e. they are not statistically different).

### Propagation of uncertainty

The uncertainty around the metric *O*_*τ*_ is propagated from the uncertainty of the KM estimator of *S*(*t*). In this section, this propagation is explicitly demonstrated.

### Uncertainty around conditional survival

This model is fitted in R using the `survival::survfit` function, which allows for the argument `conf.int` to be set to the desired confidence level (default is 0.95).

Our model presents a 95% confidence interval for the conditional probability of an individual of age *t* outliving age *τ, P*(*T* > *τ* ∣ *T* > *t*) (see black line in Figure 1). This conditional survival is calculated as described in Equation 8, where *S*(*t*) is estimated using a standard KM model. The conditional survival is calculated as a ratio of bounded quantities (Equation 8). For a ratio *R* = A/*B* with A, *B* > 0, *R* is largest when A is at its upper bound and *B* at its lower bound, and smallest when A is at its lower bound and *B* at its upper bound. Thus, for a ratio 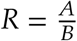, the bounds of *R* are given by:

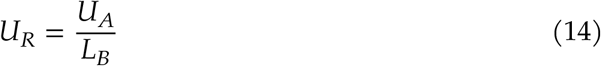

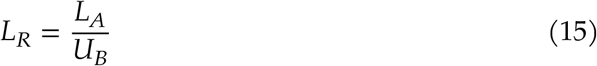

Here *P*(*U*_R_) denotes the probability that the upper bound of *R* is attained (i.e. A at its upper bound and *B* at its lower bound), and similarly *P*(*L*_R_) denotes the probability that the lower bound of *R* is attained. If the original bounds represent the *X*% CI around the original quantities A and *B*, and the two tail events are treated as independent, then 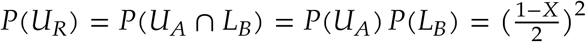, and similarly for *P*(*L*_*R*_). Thus the ratio bounds correspond to a 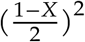 probability in each tail:

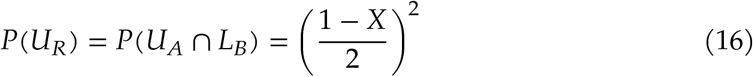

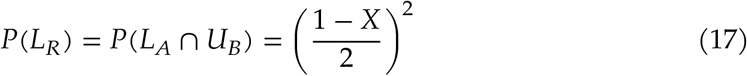

Hence, when initially computing the KM estimator, the confidence intervals must be chosen such that their ratio will yield the desired 95% CI for the conditional survival. For a 95% CI, each tail has probability 0.025; we therefore set 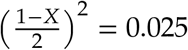 and solve for *X*:

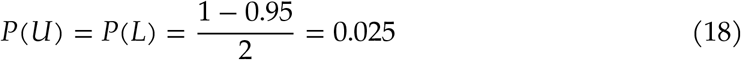

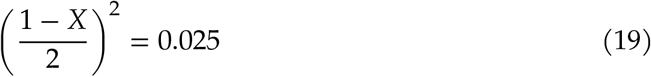

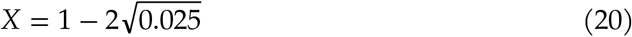

In summary, by setting `conf.int = 1-2*sqrt(0.025)` in the call to `survival::survfit`, the conditional survival bounds computed as seen in Equation 14 and Equation 15 define a 95% CI for *P*(*T* > *τ* ∣ *T* > *t*).

### Uncertainty around *O*_*τ*_

The probability of a random individual of unknown age outliving age *τ* (i.e. *O*_*τ*_) is also presented with a 95% CI. This metric is calculated as the weighted sum of the conditional survival for all possible values of *t* (see Equation 9).

The uncertainty around this metric is propagated from the uncertainty of the conditional survival alone, since A(*t*) (the age distribution) is treated like a fixed probability distribution. Because *O*_*τ*_ is a weighted sum (Equation 9) and each term is positive, the upper (lower) bound of the sum is the sum of the upper (lower) bounds of each term. Thus, the bounds of *O*_*τ*_ are given by:

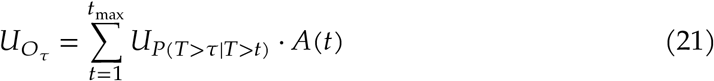

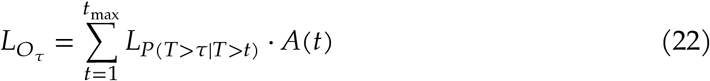

### Assumptions

The model presented here relaxes the assumptions of proportional hazards and of a parametric form for the survival distribution that are required by Cox PH and parametric survival models. However, it does make some assumptions that are important to consider when applying it to real-world data:

- **The data are sufficient to estimate the survival function** *S*(*t*) **across the range of** *t* **of interest**: All metrics used in this model are derived ultimately from *S*(*t*), so it is crucial that this function is well approximated. High sample sizes and long follow-up periods would improve the model’s precision significantly.
- **The age distribution of the population can be known**: This model assumes that the age distribution A(*t*) is a knowable (or at least, approximable) distribution. However, age structures of wild populations are rarely stable, and this has been argued theoretically [May, 1974], and confirmed experimentally [Costantino et al., 1997] and observationally [Hoy et al., 2020].

### Summary of model inputs and outputs

The model presented here takes a life table as input, which is the standard for survival analysis [Flynn, 2012]. It also requires that an approximation of the age distribution A(*t*) be provided, although this can be estimated from *S*(*t*) in the simplest case of stationary populations (see Section 3.3). The model outputs are summarised below:

- *P*(*T* > *τ* ∣ *T* > *t*), the conditional survival matrix. This is a 2-dimensional matrix where the element (*t, τ*) represents the probability that an individual of age *t* will outlive age *τ*. It is defined for all positive values of *t* and *τ* that are smaller than the largest age recorded in the study. This matrix is a function of both *t* (i.e. the age of the individual) and *τ* (i.e. the target age beyond which survival is of interest).
- *O*_*τ*_(*t*), the probability of outliving age *τ* conditional on the current age *t*. This is equivalent to the *τ*-th row of the conditional survival matrix, and is a function of *t* (i.e. the age of the individual).
- *O*(*τ*), the probability of outliving age *τ* when the age is unknown, as a function of *τ*. This function may look similar in shape to *S*(*t*), but it is fundamentally different. While *S*(*t*) describes the probability *at birth* of surviving until age *t, O*(*τ*) describes the probability *of a randomly sampled individual* of surviving until age *τ*. This is done by averaging the conditional survival matrix over the *t* dimension using the age distribution A(*t*) as a weighing function.
- *O*_*τ*_, the probability of outliving age *τ* when the age is unknown, for a particular value of *τ*. This can be understood as a specific case of *O*(*τ*), or, equivalently, as the area under the curve of *O*_*τ*_(*t*) weighed by A(*t*).
- Confidence intervals for all of the above (default is 95%, but can be customised).
- Pairwise statistical comparison of *O*_*τ*_ estimates, for any two groups and *τ* values of interest (different groups can have different values for *τ*).

### Model assessment using simulated and real-world data

To evaluate the performance and practical utility of our survival modelling framework, we conducted one simulation study and downloaded two publicly available real-world ecological datasets. We then compared our model to the two most common approaches to survival analysis (Cox’s PH model and parametric survival model).

The simulated dataset included survival data on two allopatric populations of a hypothetical species, hereafter referred to as Island A and Island B. Each island had a specific birth rate and baseline life expectancy. All individuals had a 50% chance of carrying a mutation, which affected their odds of survival in a complex way. For individuals with a low baseline life expectancy (< 15), the mutation increased their survival. For those with high baseline life expectancy (> 25), the mutation was beneficial in Island A and detrimental in Island B. Table 1 shows the exact processes and parameters used to simulate these data.

**Table 1:**
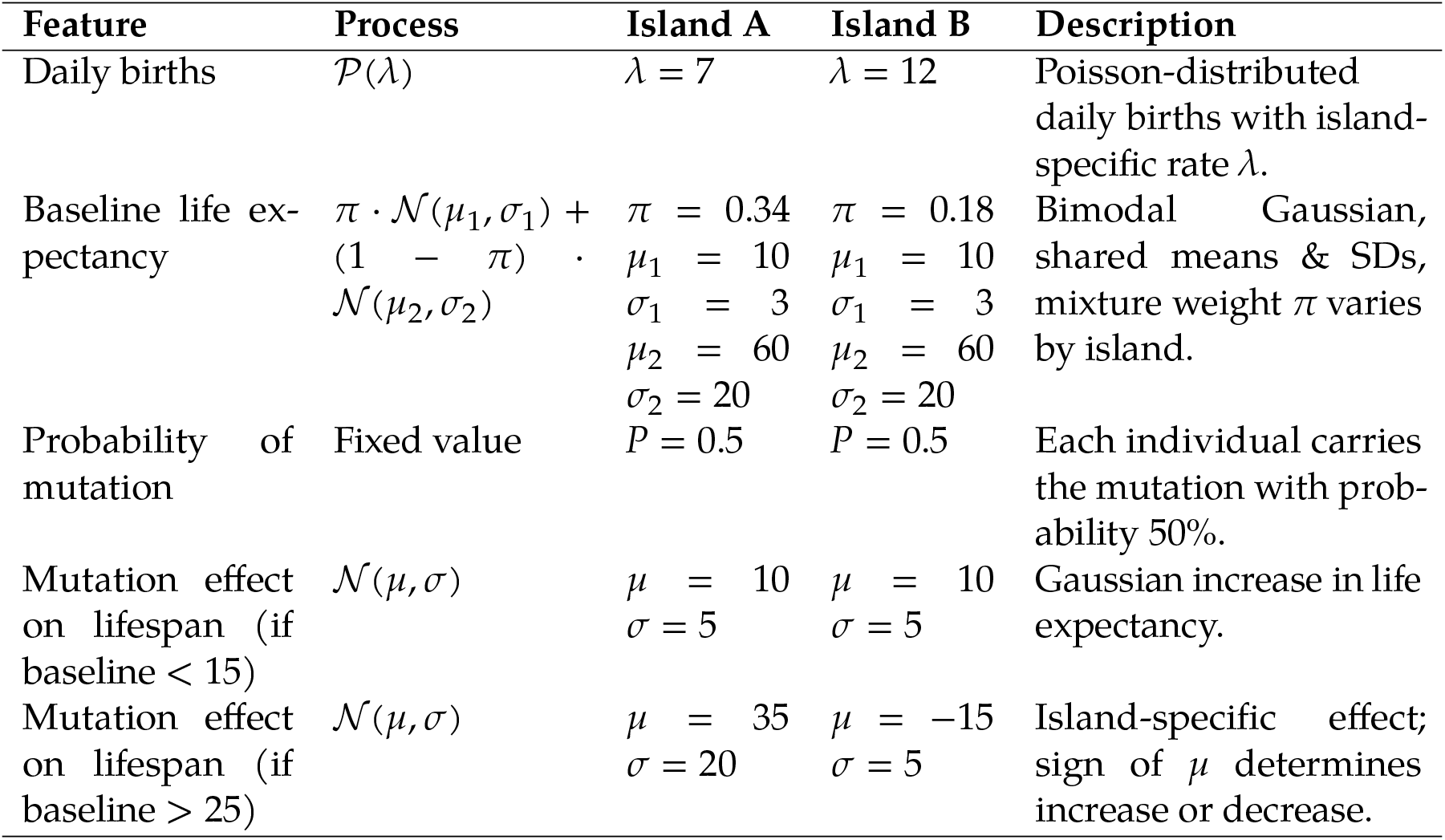
Processes and parameters used to simulate the survival dataset.

| Feature | Process | Island A | Island B | Description |
| --- | --- | --- | --- | --- |
| Daily births | $\mathcal{P}(\lambda)$ | $\lambda = 7$ | $\lambda = 12$ | Poisson-distributed daily births with island-specific rate $\lambda$. |
| Baseline life expectancy | $\pi \cdot \mathcal{N}(\mu_1, \sigma_1) + (1 - \pi) \cdot \mathcal{N}(\mu_2, \sigma_2)$ | $\pi = 0.34$<br>$\mu_1 = 10$<br>$\sigma_1 = 3$<br>$\mu_2 = 60$<br>$\sigma_2 = 20$ | $\pi = 0.18$<br>$\mu_1 = 10$<br>$\sigma_1 = 3$<br>$\mu_2 = 60$<br>$\sigma_2 = 20$ | Bimodal Gaussian, shared means & SDs, mixture weight $\pi$ varies by island. |
| Probability of mutation | Fixed value | $P = 0.5$ | $P = 0.5$ | Each individual carries the mutation with probability 50%. |
| Mutation effect on lifespan (if baseline $< 15$) | $\mathcal{N}(\mu, \sigma)$ | $\mu = 10$<br>$\sigma = 5$ | $\mu = 10$<br>$\sigma = 5$ | Gaussian increase in life expectancy. |
| Mutation effect on lifespan (if baseline $> 25$) | $\mathcal{N}(\mu, \sigma)$ | $\mu = 35$<br>$\sigma = 20$ | $\mu = -15$<br>$\sigma = 5$ | Island-specific effect; sign of $\mu$ determines increase or decrease. |

The simulated dataset consisted of 191,399 individuals. Prior to any analysis, we artificially subsampled the data to achieve realistic conditions in terms of censoring, sample size, and follow-up period.

Subsampling the dataset this way also enabled the assessment of a key assumption of our model, namely that the true population’s age distribution *A*(*t*) can be estimated from data (Figure 6). The subsampling was done as follows, and Figure 2 describes the final working dataset:

**Fig. 2:**
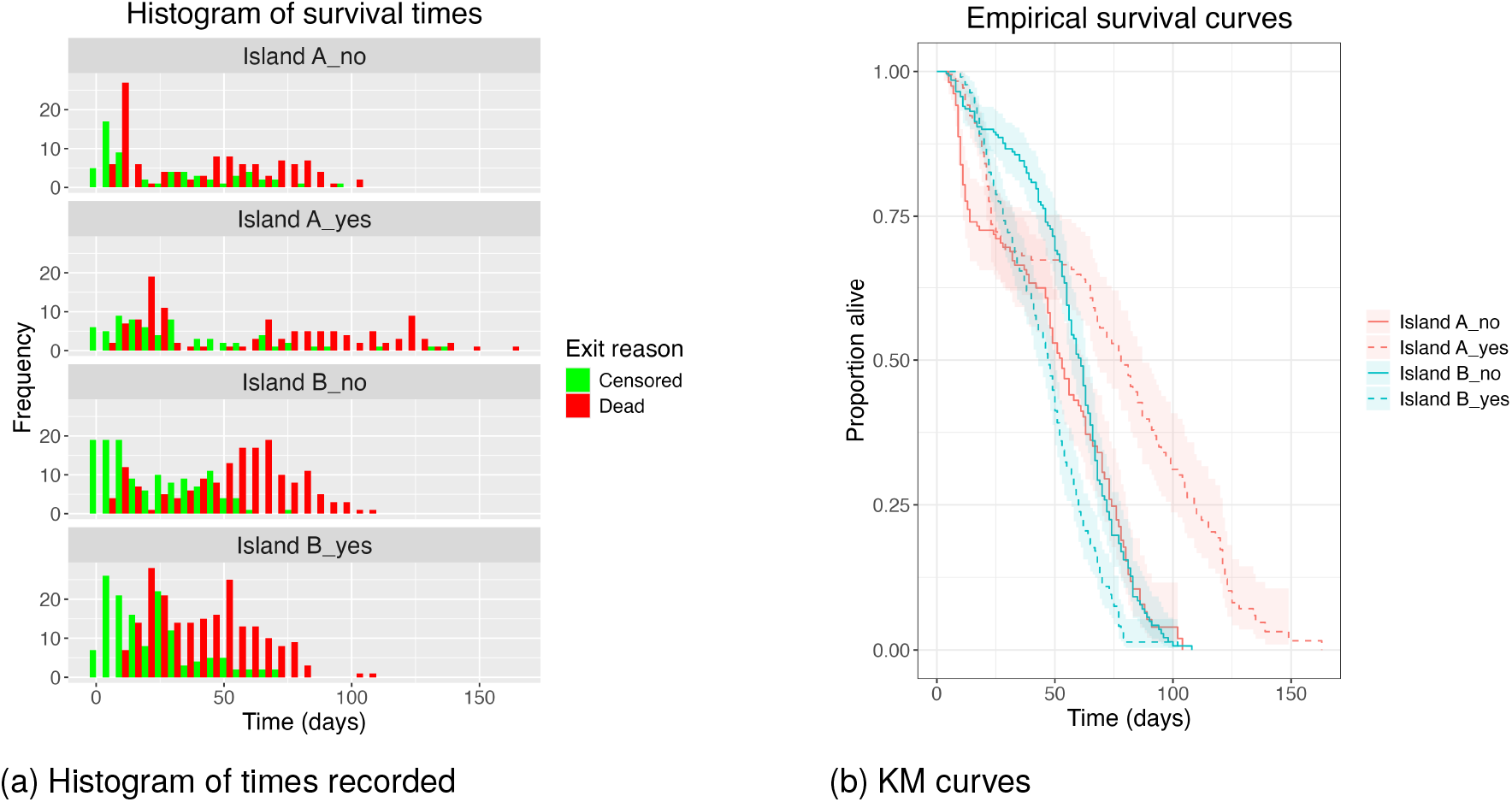
Simulation study - data description. (a) Distribution of event times in the simulation study, aggregated by population (i.e. island: A/B) and genotype (i.e. mutation status: Yes/No). The x-axis represents the age of the individuals, while the y-axis represents the number of individuals at each age. Red bars represent individuals that were followed up until death and green bars represent individuals whose follow-up ended for a different reason (right-censored). (b) Kaplan-Meier (KM) curves for each subpopulation in the simulation study. The x-axis represents time, while the y-axis represents survival probability. The data are aggregated by population (i.e. island: A/B) and genotype (i.e. mutation status: Yes/No). The 95% CI are represented by the shaded areas around each curve.

- **Censoring**: We randomly selected 40% of the individuals and truncated their recorded lifespan to a random value between 0 and their true age at death. These individuals were then marked as right-censored in the dataset.
- **Sample size**: We randomly selected 1000 individuals and discarded the rest.
- **Follow-up period**: The follow-up period was set to 200 days, which aims to represent an appropriate experimental design given the lifespan of the simulated species. Individuals that were alive at the end of the follow-up period were right-censored at their age at that time.

This simulation aimed to resemble real-world ecological survival data with credible features. In this scenario, ecologists are collecting data from two allopatric populations (e.g. bats) in a metapopulation context (e.g. an archipelago) where movement across subpopulations (e.g. islands) is possible. Of all subpopulations, only two are accessible to the ecologists (e.g. funding/logistic limitations). The hypothetical research aim is to describe the effect of a certain mutation on survival.

The two real-world case studies illustrate how Cox PH, parametric survival modelling, and our model perform when applied to real datasets. One dataset has data on the survival of *Drosophila* flies from different countries [Klepsatel et al., 2013]. The other one, on the survival of *Drosophila* flies reared at different temperatures [Gerken, 2023]. Both datasets are publicly available and were obtained from online repositories. The code used to simulate the data, and to analyse the three datasets is available in the supplementary materials (Section A) and as the R package `TAUS` (https://github.com/casasgomezuribarri/TAUS).

Cox models were fitted using the `survival::coxph` function in R. Parametric survival models were fitted using the `flexsurv` package in R. First, the most appropriate distribution was selected among all of those available in the package (i.e. Exponential, Log-Normal, Log-Logistic, Gamma, Gompertz, Weibull, Generalised Gamma, and Generalised F). This was done by fitting a null model to the whole dataset with each distribution, and then selecting the best fit through a combination of formal goodness-of-fit metrics (AIC, Log-likelihood, and KS test) and visual inspection of the models’ fit to the empirical survival, hazard, and cumulative hazard functions. The code for this is available in the supplementary materials (Section A); TAUS analyses can be reproduced using the `TAUS` R package (https://github.com/casasgomezuribarri/TAUS).

## RESULTS

### Simulation study

A simulated dataset was created as described in Section 3.8, aiming to resemble a realistic ecological scenario in terms of underlying processes, sample size, censoring, and followup period. The dataset consisted of survival data on two allopatric populations, where individuals could either carry or not a mutation that affected their survival in a complex, population-specific manner.

### Cox PH

The Cox PH model was specified with the following covariates for survival times: mutation status, island, and their interaction. As expected from the crossing KM curves (Figure 2b), the PH assumption was found to be violated (*p* << 0.05). Thus, this model was considered invalid and should not be used. However, with the purpose of output comparison, we ignore that the PH assumption is violated and proceed to interpret the model’s outputs (Table 2).

**Table 2:** Output of the Cox PH model applied to the simulated dataset.

| Variable | HR ($\exp(\beta)$) | 95% CI | p-value |
| --- | --- | --- | --- |
| Mutation (Yes) | 0.38 | 0.29 – 0.51 | $4.3 \times 10^{-11}$ |
| Habitat (Island B) | 0.84 | 0.66 – 1.1 | 0.15 |
| Mutation (Yes) × Habitat (Island B) | 4.2 | 3.0 – 6.0 | $2.2 \times 10^{-15}$ |
| Reference level: Habitat = Island A, Mutation = No |  |  |  |
| Model statistics: $n = 1000$, events = 607 | | | |
| Concordance = 0.571 (SE = 0.014) |  |  |  |
| Likelihood ratio test = 100.8 on 3 df, $p < 2 \times 10^{-16}$ | | | |
| Wald test = 90 on 3 df, $p < 2 \times 10^{-16}$ | | | |
| Score (logrank) test = 95.04 on 3 df, $p < 2 \times 10^{-16}$ | | | |

The reported Likelihood ratio, Wald, and Score test (all with *p* << 0.05) agree on the good fit of the model. However, the Concordance index reveals a predictive performance (0.57) extremely close to random (0.5) [Longato et al., 2020]. Altogether, these statistics suggest that, although the choice of explanatory variables was appropriate, the model was not able to accurately describe the survival dynamics of the system, which is consistent with the violation of the PH assumption onto which Cox PH models rely.

The hazard ratio for the variable *mutation* (*HR* = 0.38, *p* << 0.05) indicates that, at any point in time, the immediate risk of death for individuals with the mutation from the reference island (Island A) is 38% of that for individuals from the reference level (island: A, mutation: No). The output also suggests that individuals from Island B with the reference mutation status (No mutation) have an immediate risk of death that is not significantly different from that of individuals from the reference level (island: A, mutation: No; *HR* = 0.84, *p* = 0.149). Lastly, the interaction term indicates that individuals from Island B with the mutation have an immediate risk of death that is, at all times, 0.38×0.84×4.2 = 1.34 times that of individuals from the reference level (*p* << 0.05).

### Parametric survival

The Weibull distribution was found to be the best fit available for the parametric survival analysis, which prompted an Accelerated Failure Time (AFT) model [Jackson et al., 2011]. As with the Cox PH model, this was specified with the following covariates for survival times: mutation status, island, and their interaction (Table 3, Figure 3).

**Table 3:** Output of the parametric survival model (Weibull, AFT) applied to the simulated dataset.

| Variable | Estimate ($\beta$) | AFT factor ($\exp(\beta)$) | Std. error | Statistic | p-value |
| --- | --- | --- | --- | --- | --- |
| Shape ($k$) | 2.0 | NA | 0.065 | NA | NA |
| Scale ($\lambda$) | 58.0 | NA | 2.7 | NA | NA |
| Mutation (Yes) | 0.36 | 1.4 | 0.065 | 5.6 | $2.3 \times 10^{-8}$ |
| Habitat (Island B) | 0.097 | 1.1 | 0.061 | 1.6 | 0.11 |
| Mutation (Yes) × Habitat (Island B) | -0.59 | 0.88 | 0.083 | -7.1 | $1.6 \times 10^{-12}$ |

**Fig. 3:**
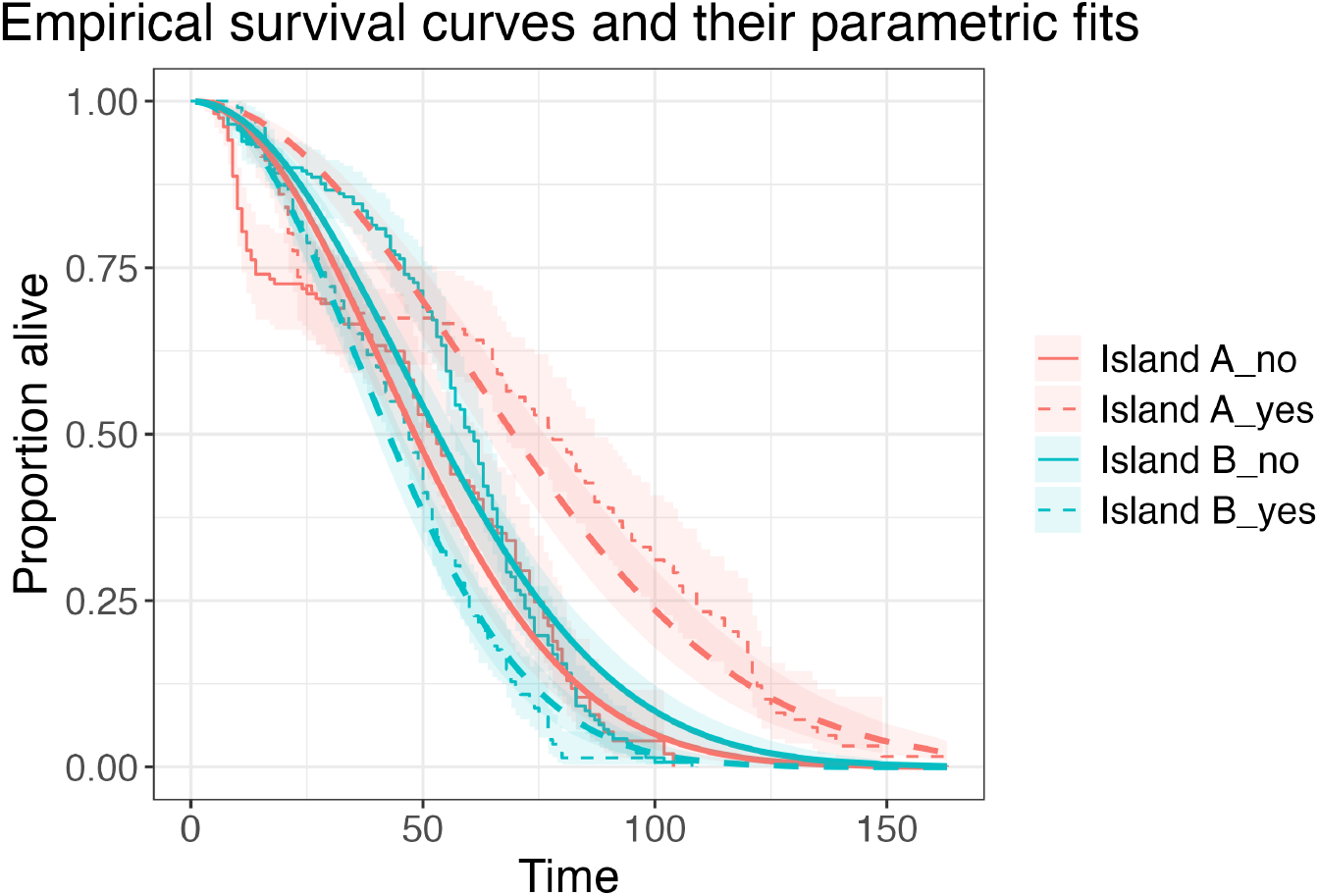
Empirical KM curves of each subpopulation in the simulation study (step curves), overlapped with their respective parametric fits (smooth curves). Data are aggregated by island (A: red, B: blue) and mutation (No: solid lines, Yes: dashed lines). Shaded regions represent the 95% confidence intervals.

The output (Table 3) lists the parameters for the baseline Weibull hazard function and how the specified covariates modify them. The shape parameter *k* ≈ 2 is common to all groups, and indicates that the risk of death increases over time in a roughly linear manner [Incerti, 2019]. The scale parameter controls the stretching of the x-axis (i.e. time), acting as a constant that sets the overall death rate (or, more precisely, the speed at which time is experienced). The coefficients reported by the model are the modifications to the scale parameter due to each explanatory variable [Jackson et al., 2011]. The coefficient of the variable *mutation* indicates that individuals from the reference island (Island A) with the mutation experience time 1.4 times slower than the baseline group (island: A, mutation: No; *p* << 0.05), meaning that their lifespan is increased by 40%. Similar to the Cox model, the effect of island alone was not significant when the mutation status was at its reference level (No mutation; *p* = 0.11). The interaction term indicates that individuals from Island B with the mutation experience time 0.88 times as fast as the baseline group, which means that their rate of death is increased by 12% (*p* << 0.05). This result is graphically presented in Figure 3.

This modelling approach provided a better description of the survival dynamics than the Cox PH model by explicitly accounting for the baseline hazard function, which removed the PH assumption [Cox and Reid, 1994]. However, the Weibull distribution was a poor fit to the empirical survival functions, which is illustrated by the general lack of shape overlap between the empirical and fitted curves, particularly at early times (Figure 3). Furthermore, the description of covariate effects on survival was limited to a single, time-invariant value per covariate (i.e. the AFT factor).

### TAUS model

The TAUS model is implemented in the R package `TAUS` (https://github.com/casas-gomezuribarri/TAUS). The previous sections demonstrate that Cox PH and parametric survival models were limited in their ability to describe the simulated system. The Cox model was invalid due to the violation of the PH assumption, and should not have been considered at all. The parametric model was valid, but its fit to the empirical data was imperfect (the shape of the fitted curves did not match the shape of the empirical curves), and its description of covariate effects on survival was limited to a stretching/compressing parameter for this imperfect fit along the time dimension. The TAUS model, on the other hand, makes no assumptions about the hazard function, and relies entirely on the empirical survival function. For each population, it computes all possible conditional survival probabilities (i.e. *P*(*T* > *τ* ∣ *T* > *t*) for all possible combinations of *τ* and *t*) and compiles them into a matrix. Figure 4 shows these as heatmaps.

**Fig. 4:**
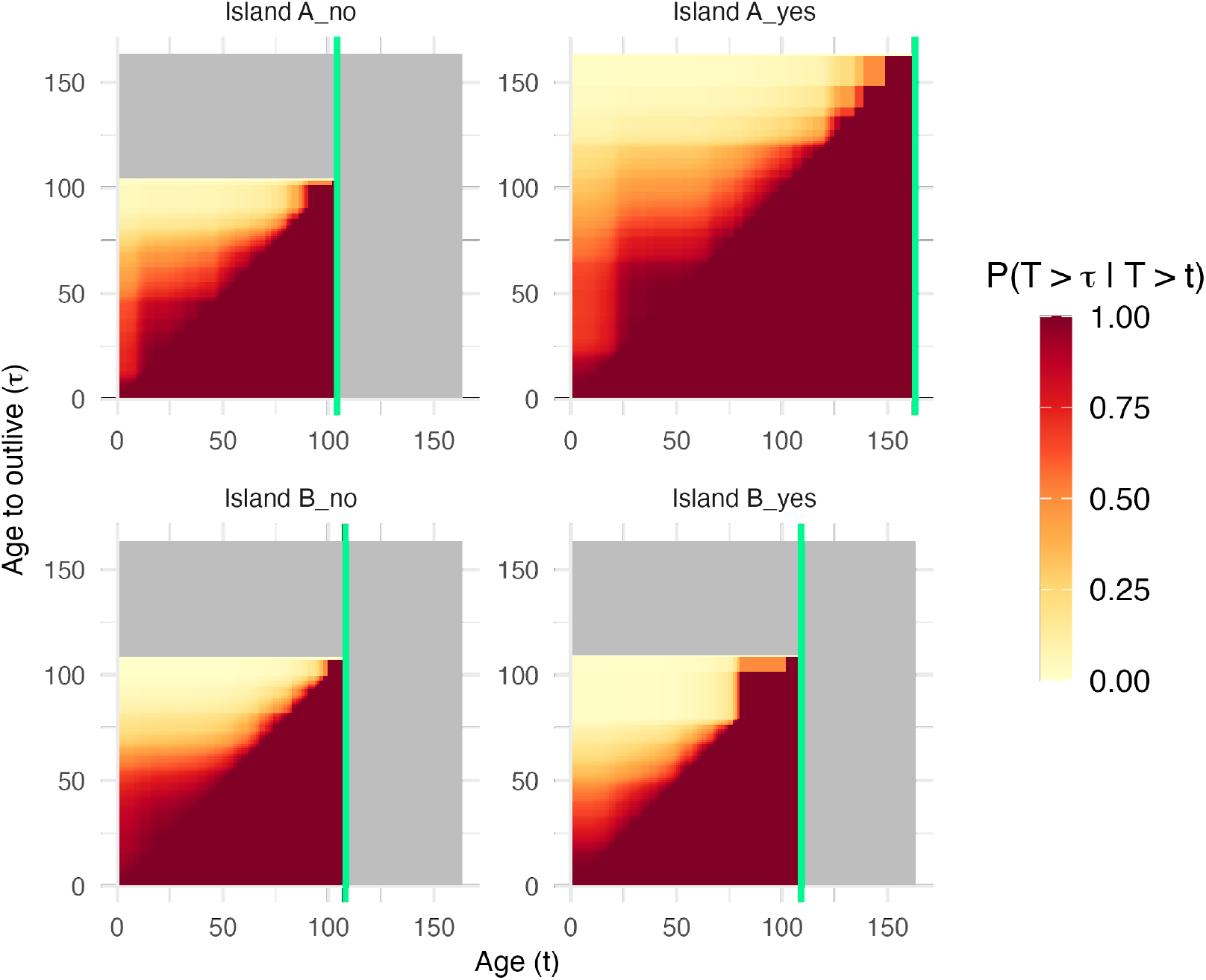
Each panel shows the conditional survival matrix for one of the four groups in the simulation study. The x-axes represent the current age of the individuals (*t*), while the y-axes represent the age beyond which survival is of interest (*τ*). The colour of each cell (*x, y*) represents the probability that an individual of age *t* = *x* will outlive age *τ* = *y*, that is, *P*(*T* > *τ* ∣ *T* > *t*). Grey cells represent coordinates (*t, τ*) for which this probability is undefined (i.e. *τ, t* ≥ maximum age observed in the group). The vertical green lines mark the time of the last death observed in each group.

From these matrices, we can compute all outputs outlined in Section 3.7 (Figure 5).

**Fig. 5:**
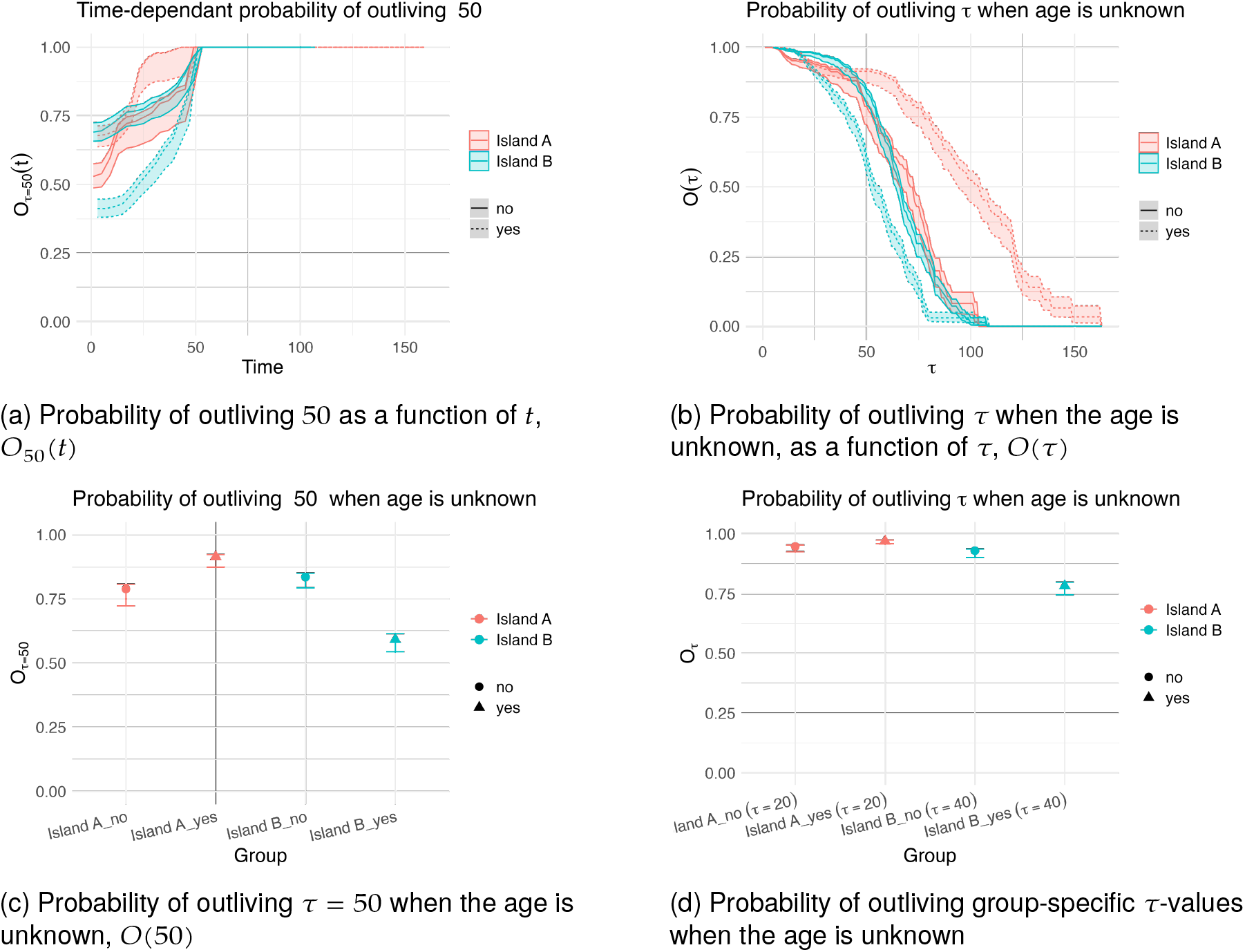
A summary of the possible outputs of the TAUS model. (a) The probability of outliving age *τ* = 50 as a function of the current age *t*. (b) The probability of outliving *τ* as a function of *τ* when the age is unknown. (c) The probability of outliving age *τ* = 50 when the age is unknown. (d) The probability of outliving group-specific ages (*τ* = 20 for island A and *τ* = 40 for island B) when the age is unknown. All panels include 95% CI limits and aggregate the data in four groups, according to island of origin (A/B) and mutation status (yes/no).

**Fig. 6:**
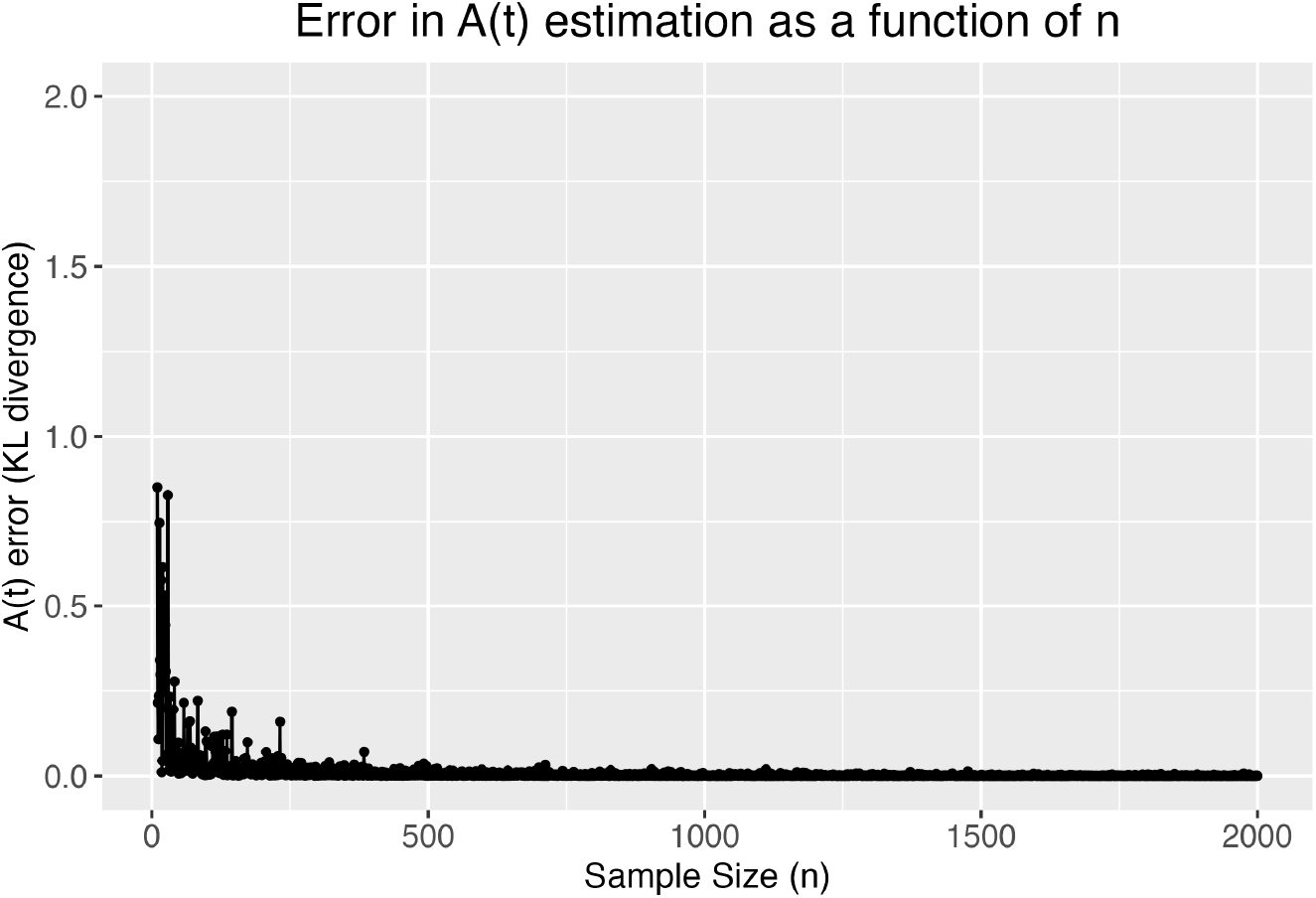
Power analysis based on the accuracy of the estimation of the true age distribution (*A*(*t*)), computed as the KL divergence between the estimated and ground truth distributions, as a function of sample size.

Figure 5a shows that individuals with a mutation from island A have a consistently higher probability of outliving the arbitrary age of *τ* = 50 than the other groups, regardless of their current age (i.e. highest value of *O*_50_(*t*), regardless of *t*). It also shows that individuals without the mutation have a similar age-dependent probability of outliving *τ* = 50 during most of their lives, regardless of their island of origin. As expected, the probability of outliving age 50 is 1 for any individual that is already 50 or older.

Figure 5b shows that this trend is true for almost any age *τ* to be outlived, not just *τ* = 50. Individuals from island A with the mutation have the highest probability of outliving virtually any age. Individuals with the mutation from island B have the highest probability of outliving young ages, but their chances of surviving beyond any age greater than *τ* ≈ 30 are the lowest in the study. Individuals without the mutation have similar chances of outliving most ages, but those from island A experience higher early life mortality.

Figure 5c shows that the probability of a randomly-sampled individual (and thus, of unknown age) from island A of outliving the age of *τ* = 50 is higher than for any other population. It also suggests that this probability is similar for individuals without the mutation, regardless of their island of origin. Lastly, a random individual with the mutation from island B would have the lowest chances of living beyond this age.

Figure 5d is similar, but in this case, the age beyond which survival is of interest is island-specific. Arbitrarily, we chose *τ* = 20 for island A and *τ* = 40 for island B, although any value of *τ* can be chosen for any of the four populations. Figure 5d shows that having the mutation on island A has a positive effect on the chances of individuals surviving beyond *τ* = 20, although this effect is non-significant (Table 4). On the other hand, on island B, the effect of the mutation on chances of survival beyond *τ* = 40 is negative and significant (Table 4).

**Table 4:** All pairwise comparisons of *O*_*τ*_ values (as shown in Figure 5d). For each group, the table shows the subpopulation, the value of *τ* (which is island-specific), and *O*_*τ*_ (probability of living beyond *τ* when age is unknown). The final column shows the p-value of the KS test comparing both groups.

| Population | $\tau$ | $O_\tau$ | Population | $\tau$ | $O_\tau$ | p-value |
| --- | --- | --- | --- | --- | --- | --- |
| Island A_no | 20 | 0.945 | Island A_yes | 20 | 0.968 | 0.159 |
| Island A_no | 20 | 0.945 | Island B_no | 40 | 0.923 | 0.785 |
| Island A_no | 20 | 0.945 | Island B_yes | 40 | 0.714 | < 0.001 |
| Island A_yes | 20 | 0.968 | Island B_no | 40 | 0.923 | 0.012 |
| Island A_yes | 20 | 0.968 | Island B_yes | 40 | 0.714 | < 0.001 |
| Island B_no | 40 | 0.923 | Island B_yes | 40 | 0.714 | < 0.001 |

Together, these figures illustrate the versatility of the output of the TAUS model. For example, inspection of Figure 5b suggests that, for individuals without the mutation —and regardless of island of origin— the probability of living beyond ages > 50 is very similar. In contrast, those from island B have slightly higher chances of surviving beyond ages < 50. It is also evident that the mutation is detrimental on island B while beneficial on island A. All of this is consistent with the processes used to simulate the data (Table 1).

Figures 5c and 5d showcase how the TAUS model output can be used to compare survival probabilities across groups for specific ages of interest, which can be different for each group. This is particularly relevant in ecological settings where the chronological age at which something (e.g. metamorphosis) happens may vary across allopatric populations due to the specific conditions of their respective environments. Crucially, the model allows these comparisons to be statistically tested. Table 4 illustrates this for the comparisons shown in Figure 5d.

The strongest assumption of the TAUS model is that the survival function *S*(*t*) (which is proportional to the age distribution A(*t*) for stable populations — no growth, no migration) is well estimated. In real-world cases, this is often impossible to assess. However, for our simulated dataset, we can compute the age distribution using different subsample sizes, and calculate the KL divergence to compare these estimates with the true age distribution (as computed with the whole dataset, *n* = 191, 399). Figure 6 shows that, beyond a sample size of *n* ≈ 1000, the estimation of the true age distribution does not improve much with larger sample sizes. For this specific case study, a sample size under *n* ≈ 500 would have likely not been sufficient.

### Example 1: Flies from different countries

The dataset analysed here (publicly available [Klepsatel et al., 2013]) contains data on fly survival, origin, fecundity, fertility, and physiology. In order to keep this example focused, some variables were removed so that only one explanatory variable was considered. After pre-processing, the dataset contained survival data on wild-caught *Drosophila* flies from three different countries (Figure 7).

**Fig. 7:**
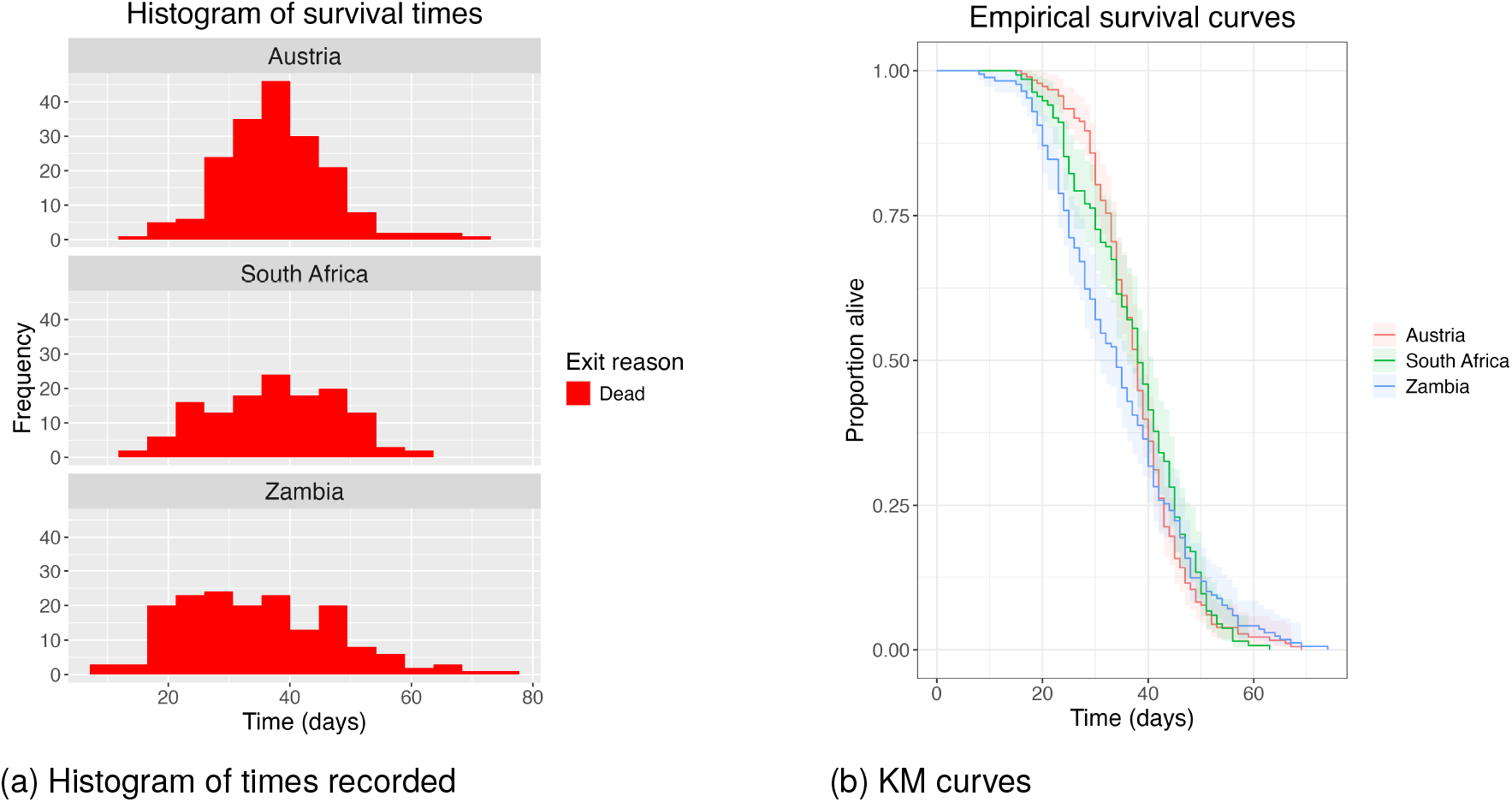
(a) Distribution of event times of the three populations in the study. The x-axis represents the age of the individuals, while the y-axis represents the number of individuals at each age. Red bars represent individuals that were followed up until death and green bars represent individuals whose follow-up ended for a different reason (censored). Note the absence of censored individuals in this study. (b) Kaplan-Meier curves for each population in the study. The x-axis represents time, while the y-axis represents survival probability.

The survival dynamics of all three populations were comparable, but with some differences. Austrian flies had the lowest mortality rates at early and late ages, but showed the highest mortality rates at mid-ages. Zambian flies had the highest mortality rate at early ages, but they also recorded the highest lifespans. South African flies lay somewhere in the middle. One of the key features of this dataset is the lack of censored individuals: all flies were observed until death (Figure 7).

### Cox PH

The Cox model was specified with country of origin as the only covariate for the survival times. The model was found to be invalid due to the violation of the PH assumption (*p* << 0.05). Again, this was expected from the crossing of the KM curves in Figure 7b.

The output of this model is reported in Table 5 for completion purposes. However, due to its invalidity, this model was not considered any further and its output will not be discussed here.

**Table 5:**
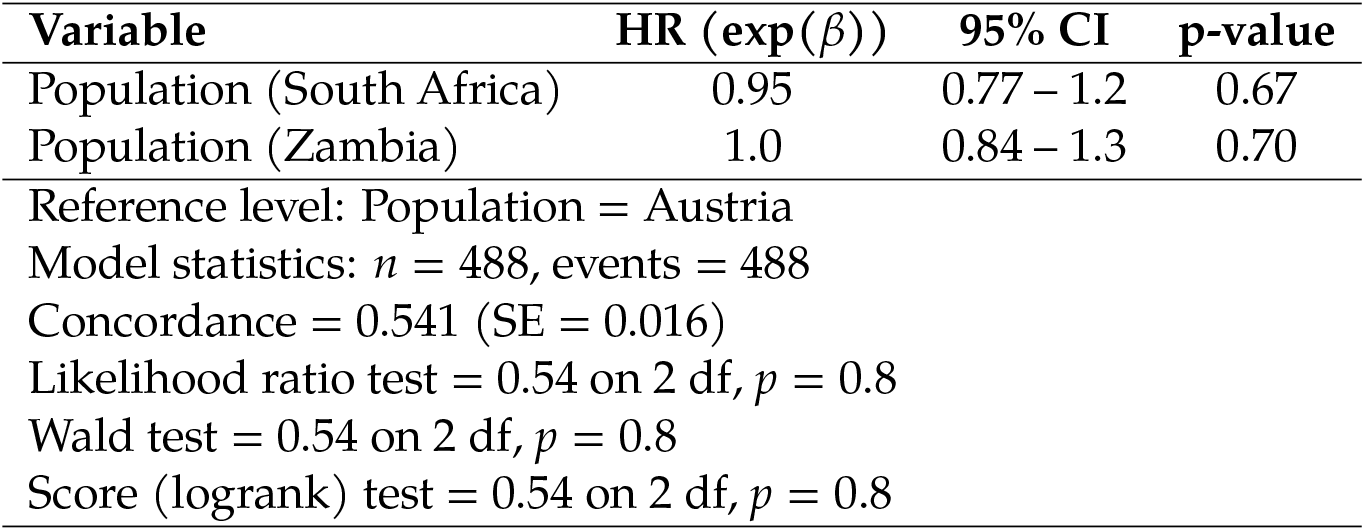
Output of the Cox PH model applied to the dataset from Example 1: Flies from different countries.

### Parametric survival

The parametric survival model was fitted using the Weibull distribution, which again prompted an AFT model. It was specified with country of origin as the sole explanatory variable. The model estimated survival to be identical across populations (Table 6, Figure 8).

**Table 6:** Output of the parametric survival model (Weibull, AFT) applied to the dataset from Example 1: Flies from different countries.

| Variable | Estimate ($\beta$) | AFT factor ($\exp(\beta)$) | Std. error | Statistic | p-value |
| --- | --- | --- | --- | --- | --- |
| Shape ($k$) | 3.7 | NA | 0.13 | NA | NA |
| Scale ($\lambda$) | 41 | NA | 0.83 | NA | NA |
| Population (South Africa) | 0.0042 | 1.0 | 0.031 | 0.14 | 0.89 |
| Population (Zambia) | -0.0030 | 1.0 | 0.029 | -0.10 | 0.91 |

**Fig. 8:**
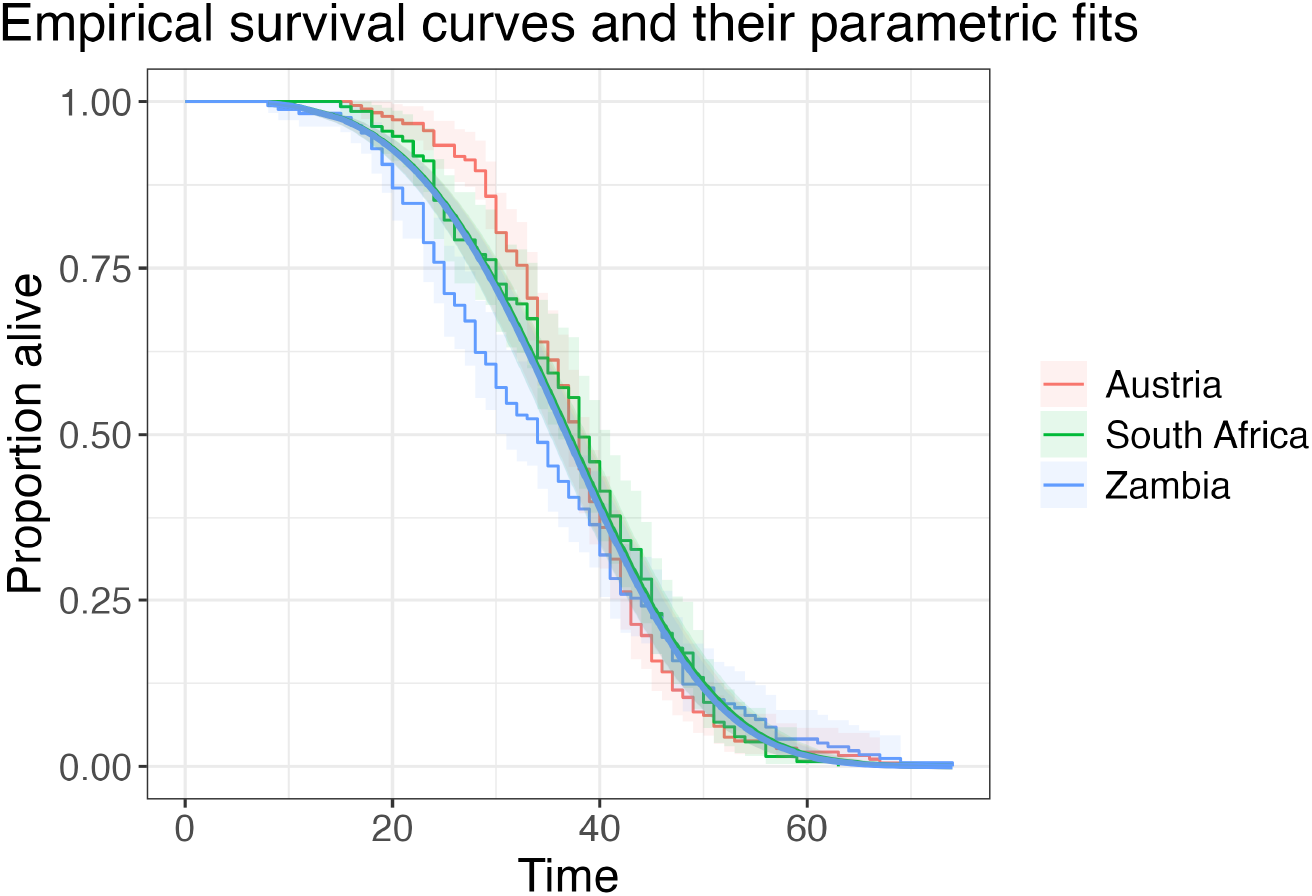
KM curves of each population in the study (step curves), overlapped with their respective parametric fits (smooth curves). Confidence intervals (95%) are shown as shaded regions.

### TAUS model

Figure 9 shows the conditional survival matrices for this dataset (aggregated by country), from which all outputs of the TAUS model are calculated.

**Fig. 9:**
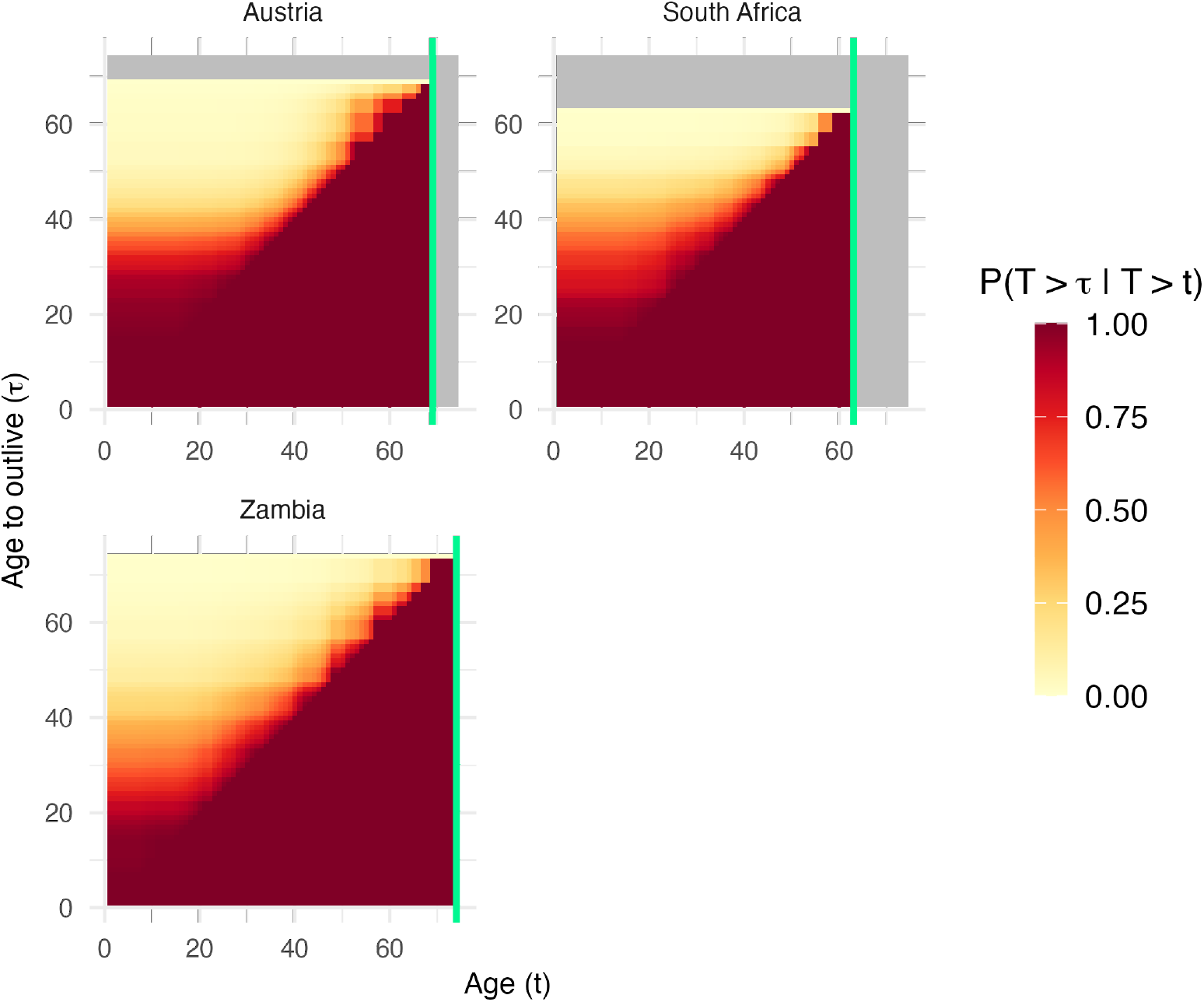
Each panel shows the conditional survival matrix for one of the groups in the study. The x-axes represent the current age of the individuals (*t*), while the y-axes represent the age beyond which survival is of interest (*τ*). The colour of each cell (*x, y*) represents the probability that an individual of age *t* = *x* will outlive age *τ* = *y*, that is, *P*(*T* > *τ* ∣ *T* > *t*). Grey cells represent coordinates (*t, τ*) for which this probability is undefined (i.e. *τ, t* ≥ maximum age observed in the group). The vertical green lines mark the time of the last death observed in each group.

From the survival matrices, we can compute the *O*_*τ*_ curves (i.e. the probability of living beyond the age of *τ* for a randomly sampled individual of unknown age, as a function of *τ* — Figure 10a). These show that Zambian flies are less likely to outlive young ages (< 40 days), and more likely to outlive old ages (> 50 days) than the other groups. This suggests that Zambian flies experience a higher early-life mortality rate, but that past a certain age, their mortality is lower than in the other populations, which is consistent with their true survival dynamics (Figure 7a).

**Fig. 10:**
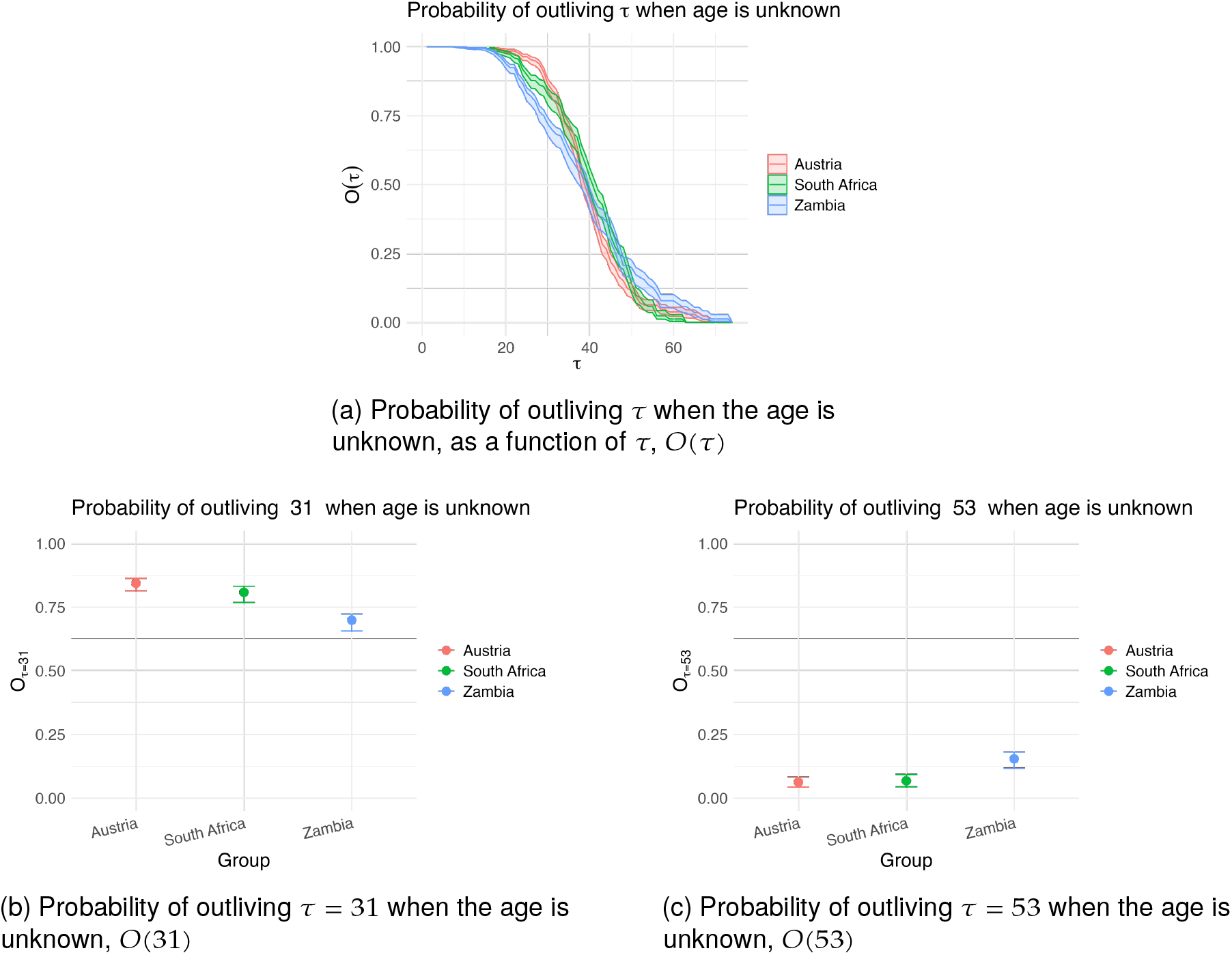
Outputs of the TAUS model describing the survival dynamics of flies from Austria (red), South Africa (blue), and Zambia (green). (a) The probability of outliving age *τ* as a function of *τ* of a randomly sampled individual of unknown age. (b) The probability of outliving age *τ* = 31 of a randomly sampled individual of unknown age. (c) The probability of outliving age *τ* = 53 of a randomly sampled individual of unknown age. Shaded region (a) and error bars (b and c) show the 95% CI bounds.

Indeed, Figures 10b and 10c show that, although a randomly-sampled fly from the Zambian population is less likely to survive age 31 than a counterpart from South Africa or Austria, it is more likely to survive age 53. The statistical significance of these comparisons is shown in Table 7. The other models (Cox PH and parametric) were not able to capture these differences (Table 5, Table 6, Figure 8).

**Table 7:** Pairwise comparisons of *O*(*τ*) for two values of *τ*.

| Comparison | | $O(\tau = 31)$ | | $O(\tau = 53)$ | |
| --- | --- | --- | --- | --- | --- |
| Group 1 | Group 2 | p-val | Reject $H_0$? | p-val | Reject $H_0$? |
| Austria | South Africa | 0.788 | No | 1.000 | No |
| Austria | Zambia | 0.002 | Yes | 0.015 | Yes |
| South Africa | Zambia | 0.012 | Yes | 0.051 | No? |

### Example 2: Flies at different temperatures

The dataset used in this section (which is publicly available [Gerken, 2023]) contains data on fly survival, strain, and rearing temperature. Similar to the previous example, some variables were removed to keep the case study focused on the analysis of how a single variable affected survival. After pre-processing, the dataset contained survival data on *Drosophila* flies reared at different temperatures. Graphical representation of the observed events (deaths and censoring, Figure 11a), and the corresponding KM curves (Figure 11b), show that higher temperatures lead to higher mortality at younger ages, although individuals with extremely long lifespans (> 200 days) were present in all groups.

**Fig. 11:**
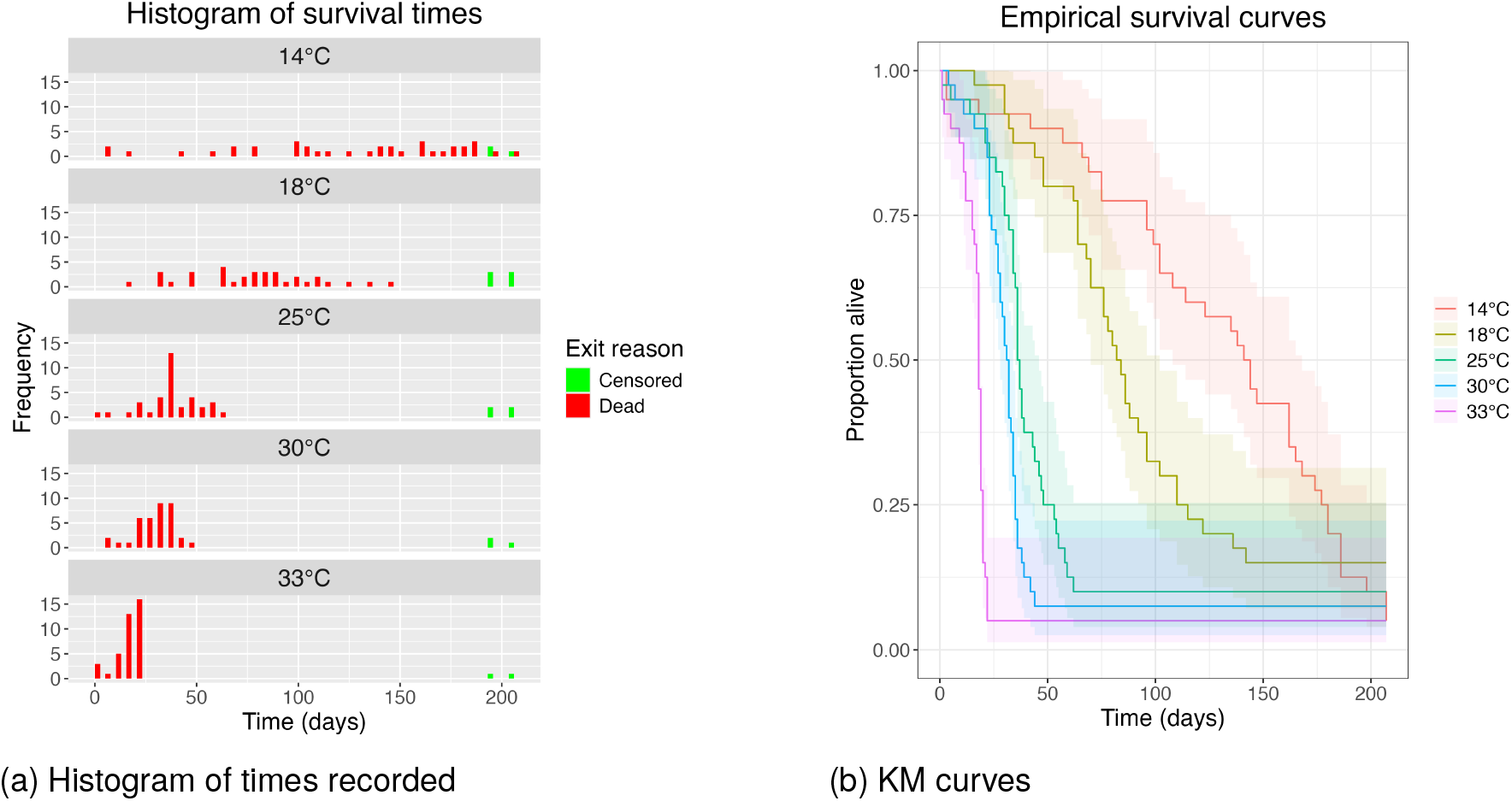
Example 2 - data description. (a) Distribution of event times of the five populations in the study. The x-axis represents the age of the individuals, while the y-axis represents the number of individuals at each age. Red bars represent individuals that were followed up until death and green bars represent individuals whose follow-up ended for a different reason (censored), in this case due to the end of the experiment. (b) Kaplan-Meier curves for each population in the study. The x-axis represents time, while the y-axis represents survival probability at birth.

### Cox PH

A Cox model was fitted to these data with temperature as the only covariate for survival times. The model was found, again, to be invalid due to the violation of the PH assumption (*p* << 0.05), and thus its output was not further investigated. However, and for completion and comparison purposes, it is reported in Table 8.

**Table 8:** Output of the Cox PH model applied to the dataset from Example 2: Flies at different temperatures.

| Variable | HR ($\exp(\beta)$) | 95% CI | p-value |
| --- | --- | --- | --- |
| Temperature (18°C) | 1.2 | 0.73 – 1.9 | 0.50 |
| Temperature (25°C) | 2.2 | 1.4 – 3.6 | $8.3 \times 10^{-4}$ |
| Temperature (30°C) | 3.2 | 2.0 – 5.2 | $1.1 \times 10^{-6}$ |
| Temperature (33°C) | 7.9 | 4.9 – 13.0 | $< 2.0 \times 10^{-16}$ |
| Reference level: Temperature = 14°C |  |  |  |
| Model statistics: $n = 200$, events = 182 | | | |
| Concordance = 0.772 (SE = 0.02) |  |  |  |
| Likelihood ratio test = 78.9 on 4 df, $p = 3.0 \times 10^{-16}$ | | | |
| Wald test = 89.5 on 2 df, $p < 2.0 \times 10^{-16}$ | | | |
| Score (logrank) test = 104.9 on 2 df, $p < 2.0 \times 10^{-16}$ | | | |

### Parametric survival

The most appropriate distribution for the parametric modelling of these survival data was the Log-logistic, which captured the most obvious general trend in the effect of temperature on *Drosophila* survival — that is, that higher temperatures increase early-life mortality (Figure 12). However, the model fails to fully capture the early-life survival dynamics, and suggests an artificial temperature-driven change in the proportion of extremely long-lived individuals. Additionally, it offers arguably poor estimates of the slopes of some of the empirical survival curves (Figure 12).

**Fig. 12:**
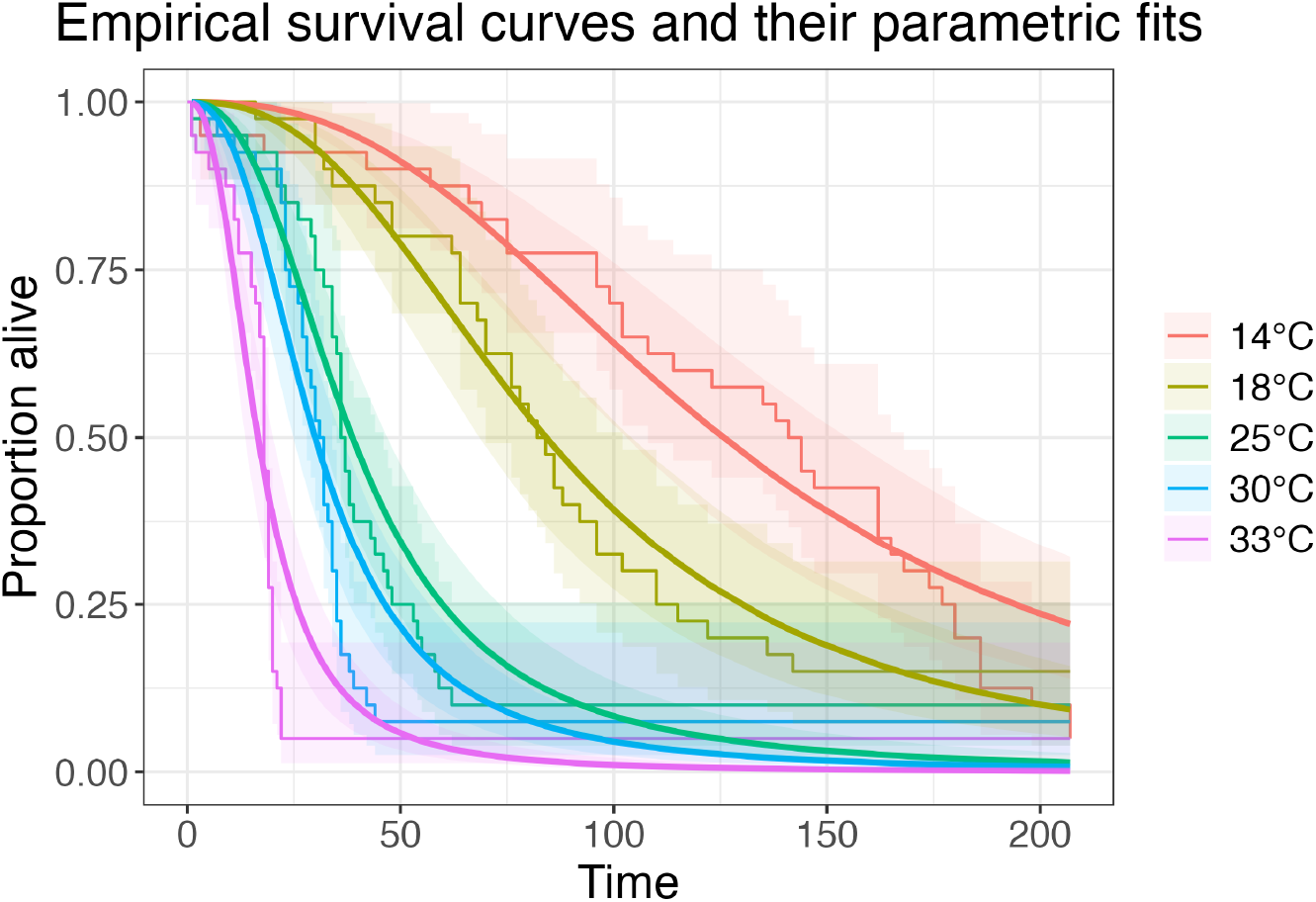
KM curves of each population in the study (step curves), overlapped with their respective parametric fits (smooth curves).

Fitting a Log-logistic model to survival data computes an Accelerated Failure Time (AFT) model, which models survival effects of covariates as dilations or contractions of the (log-) time dimension [Jackson et al., 2011]. In the case of the Log-logistic model, this is achieved by estimating the change in the scale parameter (*α*, the median of the distribution, or the age at which 50% of the population has died) as a linear function of the explanatory variables. Thus, fitting a Log-logistic parametric model to these data gives us information on how much faster individuals experience the passage of time under different rearing temperatures (Table 9, Figure 12). However, the effect of rearing temperature on *Drosophila* survival seems more complex than a simple time dilation/contraction: although the majority of the population appears to experience such effect, the proportion of extremely long-lived individuals seems to be irresponsive to their rearing temperature (Figure 11b).

**Table 9:** Output of parametric survival model fitted to the data in Example 2. The table shows the effect on the Log-logistic’s distribution location parameter (the scale parameter) of each temperature group against the reference group (14^°^C). This table is a modified version of the output that can be obtained calling the R function `flexsurv::tidy()` on the fitted model object.

| Variable | Estimate($\beta$) | $\exp(\beta)$ | Interpretation | p-val |
| --- | --- | --- | --- | --- |
| shape | 2.53 | NA | NA | NA |
| scale | 126.0 | NA | NA | NA |
| temp18°C | -0.401 | 0.669 | Survival time is 67% of baseline | 0.006 |
| temp25°C | -1.18 | 0.307 | Survival time is 31% of baseline | <0.001 |
| temp30°C | -1.43 | 0.240 | Survival time is 24% of baseline | <0.001 |
| temp33°C | -2.03 | 0.131 | Survival time is 13% of baseline | <0.001 |

This could cause misleading conclusions in certain ecological contexts. For example, in the context of R-selected species such as *Drosophila*, individuals with the longest lifespans are arguably the most relevant for population persistence in the wild, since they will undergo the most gonotrophic cycles and thus contribute the most to population growth. In an epidemiological context, extremely long-lived individuals are also likely to become superspreaders and thus contribute disproportionately to disease transmission. In both cases, a survival model that fails to capture the effects of the covariates of interest on the proportion of extremely long-lived individuals can lead to conclusions that deviate from reality in ecologically meaningful ways. This is the case of this Log-logistic parametric model, which predicts that warmer temperatures will result in a lower proportion of individuals living extremely long lives, despite this not being supported by the empirical survival curves (Figure 12).

### TAUS model

These empirical data differ from the previous example in that, for most groups, the last death recorded was a long time before the end of the follow-up period (Figure 11a). This can introduce strong survivorship bias: when most of the population dies at early ages, the longest-lived individuals account for a large proportion of the population, increasing their odds of being (randomly) sampled. This leads to the counterintuitive situation in which the probability of a randomly-sampled individual outliving old ages is highest when early mortality is also highest. The extent of the survivorship bias on this dataset can be appreciated in the conditional survival matrices (Figure 13).

**Fig. 13:**
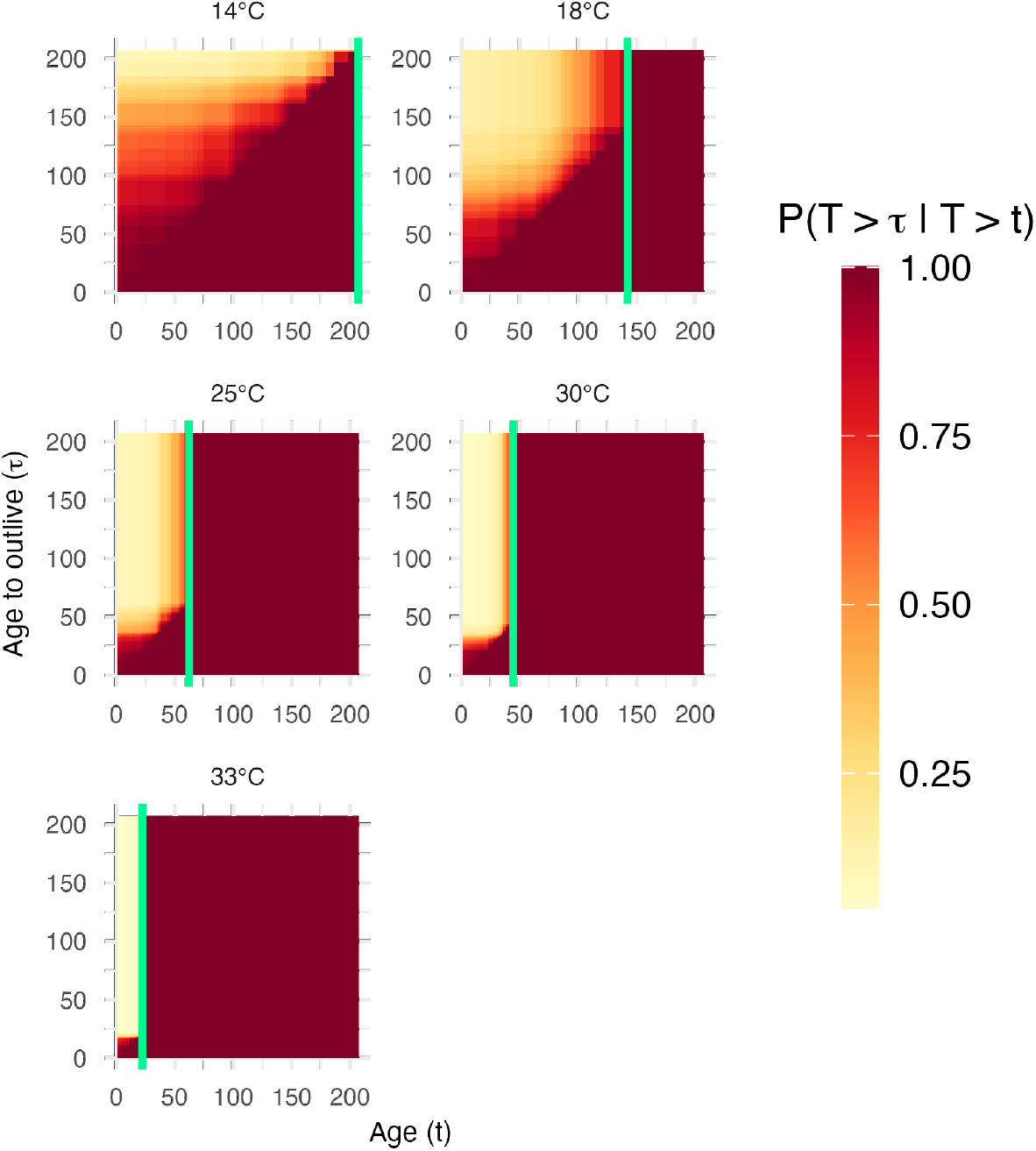
Each panel shows the conditional survival matrix for one of the groups. The x-axes represent the current age of the individuals (*t*), while the y-axes represent the age beyond which survival is of interest (*τ*). The colour of each cell (*x, y*) represents the probability that an individual of age *t* = *x* will outlive age *τ* = *y*, that is, *P*(*T* > *τ* ∣ *T* > *t*). Grey cells represent coordinates (*t, τ*) for which this probability is undefined (i.e. *τ, t* ≥ maximum age observed in the group). The vertical green lines mark the time of the last death observed in each group.

The consideration of the effects of the survivorship bias by the TAUS model is propagated from the conditional survival matrices. The *O*(*τ*) curves (i.e. the probability of a randomly sampled individual outliving age *τ* as a function of *τ*) show this very clearly (Figure 14a). Let us consider the extreme case of the 33^°^C group: most individuals died before age *τ* ≈ 25, but the remaining minority outlived the duration of the experiment, all of them reaching age *τ* > 200 (Figure 11a). This means that the probability of randomly sampling one of these extremely long-lived individuals is disproportionately high. As a result, individuals from this group have the highest *O*(*τ*) values for high values of *τ* (Figure 14a).

**Fig. 14:**
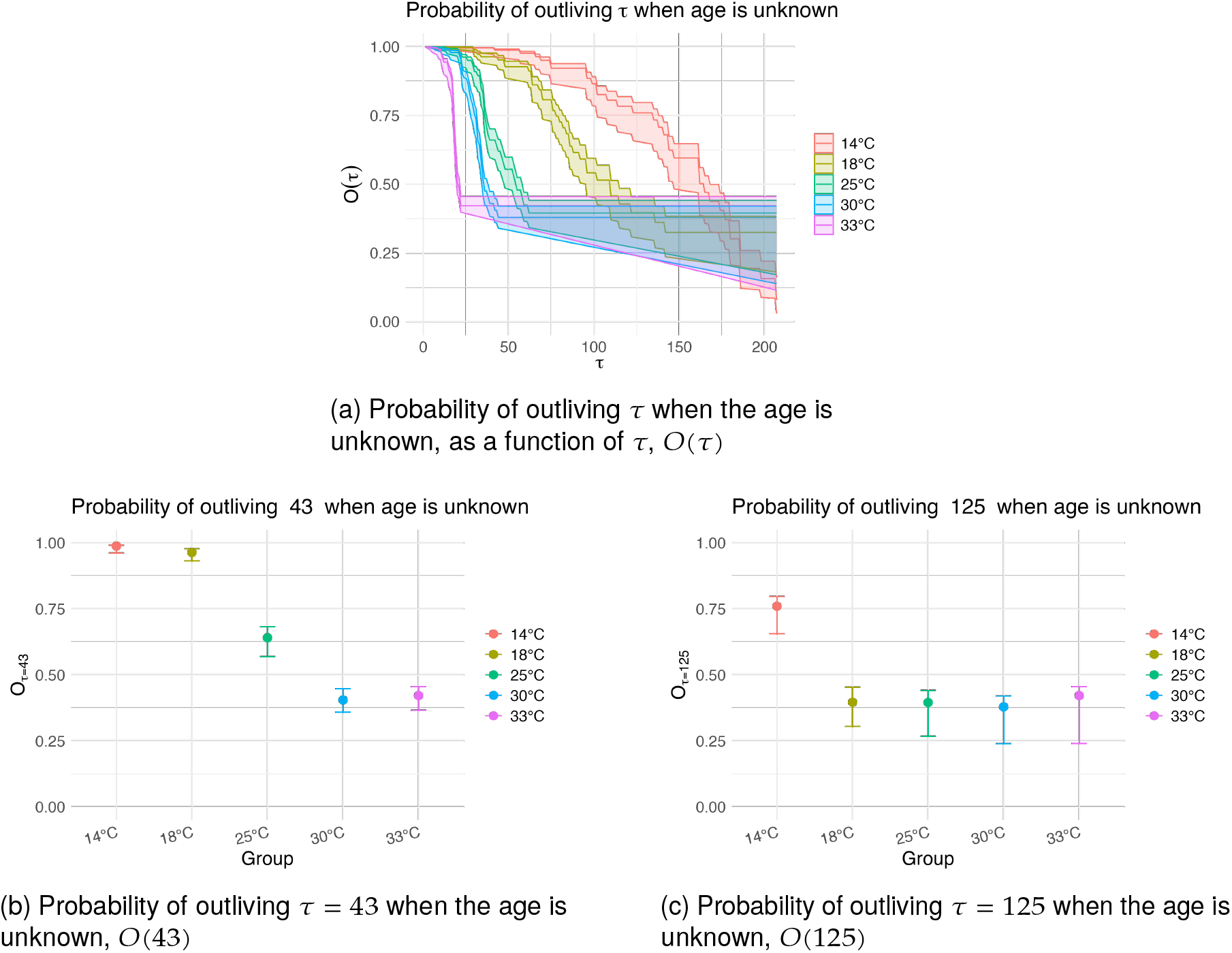
Outputs of the TAUS model describing the survival dynamics of flies reared at different temperatures. (a) The probability of outliving age *τ* as a function of *τ* of a randomly sampled individual of unknown age. (b) The probability of outliving age *τ* = 43 of a randomly sampled individual of unknown age. (c) The probability of outliving age *τ* = 125 of a randomly sampled individual of unknown age. Shaded region (a) and error bars (b and c) show the 95% CI bounds.

Thanks to the integration of the survivorship bias, the TAUS model is able to capture the general survival dynamics of the system: rearing temperatures impact early life survival (e.g. *τ* = 43, Figure 14b, Table 10), but do not affect the proportion of extremely long-lived individuals (e.g. *τ* = 125, Figure 14c, Table 10); as suggested by the histogram of event times (Figure 11a). In contrast, Cox and parametric models did not capture this (Tables 8, 9; Figure 12).

**Table 10:**
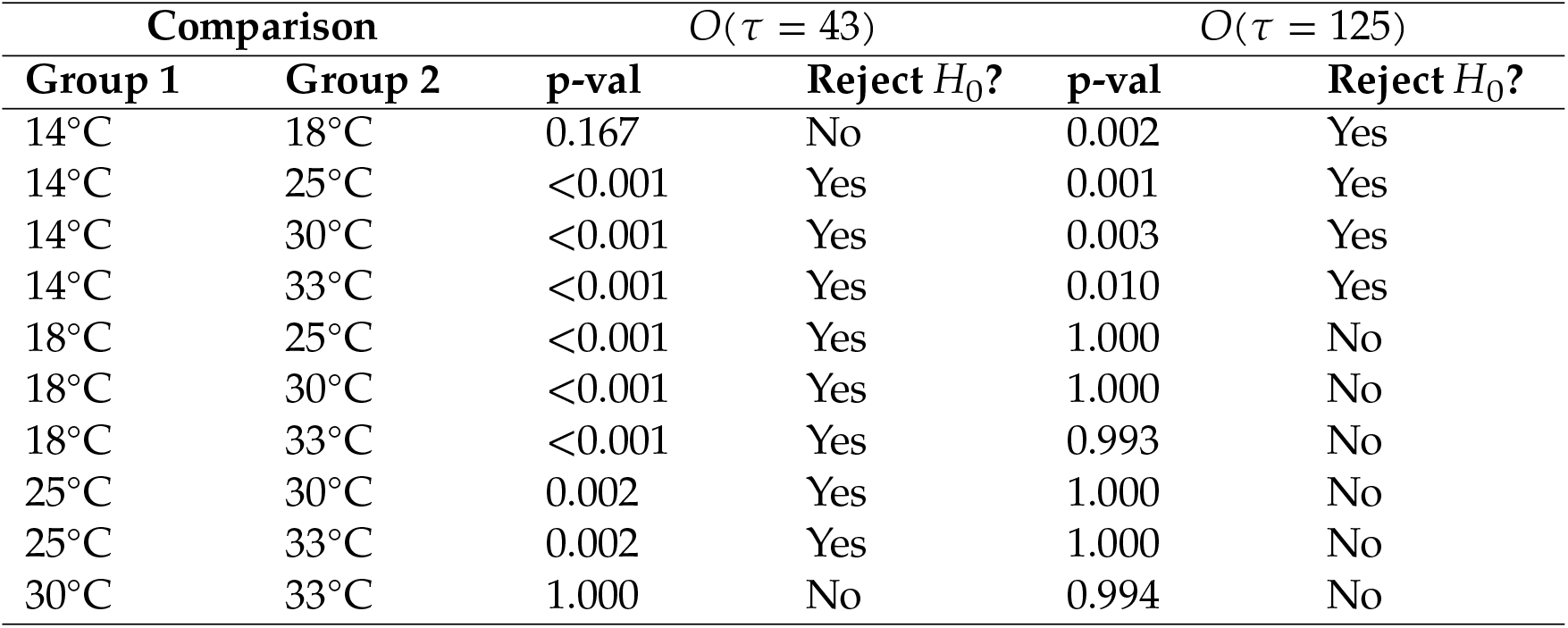
Pairwise comparisons of *O*(*τ*)

## DISCUSSION

We have developed a new method for survival analysis, TAUS, and demonstrated that it may often be better suited to ecological survival data than the most commonly used approaches, Cox PH and parametric survival models. The reason for this is two-fold. First, TAUS is free from the modelling constraints inherent to standard survival models. Second, its output is more informative, enabling detailed examination of age-specific survival probabilities.

The TAUS model does not assume proportional hazards nor does it parameterise the baseline survival function, which are common issues when applying Cox PH and parametric survival models to ecological datasets. Indeed, the PH assumption did not hold for any of the examples discussed here (simulated or real). The parametric assumption (i.e. that there exists a distribution that describes the fundamental survival dynamics of the population(s) under study) is challenging to confirm or reject beyond visual inspection of the fit under consideration. Accepting this limitation of assumption interrogation, the best fit available was selected for each example.

Noteworthily, the best fit for the two real-world examples (which included data from the same genus *Drosophila*) were different (Weibull for Example 1, Log-logistic for Example 2). If a parametric distribution existed that fundamentally described the survival dynamics of *Drosophila*, it would be expected that this distribution be the same across experiments. If the opposite is to be accepted as true —that is, that different experiments can yield different best-fitting distributions— this would suggest that no such universal distribution exists. In that case, choosing a parametric instance to describe and compare survival across experimental groups becomes somewhat arbitrary and overfitting to the specific case at hand. This is not to imply that the analysis is necessarily invalid, but rather that its generalisability is limited.

The output of the TAUS model is also more informative than the outputs of the Cox and parametric models, allowing for a richer analysis of survival data, as detailed in Section 3.7. Cox and parametric models, when epistemologically justifiable, offer relative outputs of one of two types: Proportional Hazards (PH) or Accelerated Failure Time (AFT). PH models estimate the hazard ratio (HR) between groups, which can be interpreted as ‘the immediate risk of death in population A is *HR* times that of population B’. The AFT factor, on the other hand, represents a dilation/contraction constant for the time dimension. Thus, its interpretation is of the type ‘Population A ages *AFTf actor* times as fast as population B’. Both interpretations reduce survival to a single factor change, either in risk or in time. This can be, and certainly is, useful in some contexts; but it can also miss important aspects of the survival dynamics. In contrast, the TAUS model returns the probability of outliving *τ*, where *τ* is some specific age over which survival is of interest. It also provides a framework for the statistical comparison of this probability across populations. Crucially, the value of *τ* can be chosen to be different for each population under analysis. This makes the model especially relevant to ecological systems, where the populations under comparison often have different ageing rates, and therefore the meaning of *τ* in terms of biological age can differ across them.

The probability of outliving *τ* is initially calculated as conditional on the current age *t* of the individual, allowing for fine analysis of age-dependent survival probabilities whenever appropriate. However, the model also allows the collapsing of these probabilities over the *t* dimension, effectively producing a metric for the probability of outliving *τ* when the age of the individual is unknown. This is especially useful in ecological settings, where the age of sampled wild individuals can be challenging to ascertain.

When plotting the *O*(*τ*) curves (Figures 5b, 10a, and 14a), the result can look similar to survival curves, but it is fundamentally different. While survival curves show the probability *at birth* of surviving beyond *τ*, the *O*(*τ*) curves show this probability *at the age of sampling*. As the probability of sampling older individuals increases (for example, in the presence of high early-life mortality), the *O*(*τ*) curves diverge further from the survival curves (Figure 14a). On the other hand, this divergence shrinks in cases where individuals are most likely to be sampled at young ages. This divergence is entirely due to the survivorship bias, which is the well-known logical error of concentrating analysis on samples that have passed through a selection process (in this case, being alive at the time of sampling). Despite its semantic negative connotations due to words like *bias* and *error*, this is a desirable feature in survival analysis: we are only interested in the survival of individuals that are alive.

As mentioned in Section 3.6, the TAUS model relies heavily on a good description of the age distribution of the population of interest. The simulation study presented here enabled the assessment of this assumption, but this is rarely possible in real-world cases. The age distribution of a biological system is usually subject to continuous and sometimes radical change. As a result, when applying this model, reasonable effort should be put into describing the relevant age distributions as accurately as possible. In addition, the collection of data on age-at-death for the longest lived individuals greatly facilitates the appropriate integration of the survivorship bias.

Survival analysis has traditionally been designed to investigate clinical data, where sample size is a common limitation, but information about each point in the sample is abundant. In ecological datasets the case is often the opposite: acquiring large sample sizes tends to be easier than ensuring a wealth of information about each point. In some cases, even the age of the individuals is unknown. The TAUS model is designed to be fit for purpose in this context, enabling the analysis of survival data with large sample sizes and little information per individual, and facilitating lifespan predictions on new data where the age of individuals is unknown. This is only possible because the metric for lifespan used is the probability of outliving a specific age, rather than the typical survival time (e.g. median), which is often the case.

In summary, the TAUS framework offers a flexible and ecologically grounded alternative to standard survival models by avoiding assumptions that are difficult to justify in natural systems and by producing absolute, biologically meaningful survival estimates. By aligning the statistical output with the realities of ecological processes and data collection, TAUS opens new opportunities for comparing survival across populations. We anticipate that this approach will complement existing methods and provide researchers with a principled tool for studying survival in complex ecological settings.

## Acknowledgements

The authors thank Prof Jason Matthiopoulos, Dr Mafalda Viana, and Prof Dan Haydon for their comments on early drafts of the manuscript and general intellectual engagement with the project.

## CODE AVAILABILITY

The code used to simulate the data and to analyse the datasets is available in the supplementary materials and as the R package `TAUS` at https://github.com/casasgomezuribarri/TAUS (branch `package`).

## Author contributions

**Iván Casas Gómez-Uribarri:** Conceptualisation, Data Curation, Formal Analysis, Investigation, Methodology, Software, Validation, Visualisation, Writing - original draft, Writing - review & editing; **Simon A. Babayan:** Conceptualisation, Funding Acquisition, Methodology, Resources, Software, Supervision, Writing - review & editing; **Fredros Okumu:** Conceptualisation, Funding Acquisition, Methodology, Supervision, Writing - review & editing; **Francesco Baldini:** Conceptualisation, Funding Acquisition, Methodology, Project Administration, Resources, Supervision, Writing - review & editing; **Mauro Pazmiño Betancourth:** Conceptualisation, Investigation, Methodology, Resources, Supervision, Writing - review & editing.

## Data availability statement

The data and code necessary to reproduce our analysis are available at (https://github.com/casasgomezuribarri/TAUS)

